# Exponential growth and neuronal yield in the vertebrate developing retina are mediated by Hedgehog signaling

**DOI:** 10.1101/2024.04.09.588672

**Authors:** Diego Pérez-Dones, Mario Ledesma-Terrón, Diego Mazo-Durán, Gemma Navarro-Martinez, David G. Míguez

**Author notes:** Equally contributing authors (Diego Pérez-Dones, Mario Ledesma-Terrón). Corresponding author: David G. Míguez, To whom correspondence should be addressed.

## Abstract

The balance between self-renewal and terminal differentiation of neural progenitors is regulated by a complex interplay of signaling pathways that set the spatial and temporal cues that ultimately shape and organize neurogenic tissues. In this contribution, we combine *in toto* experiments with three-dimensional image analysis, computational modeling, and theoretical tools to provide a quantitative characterization of the dynamics of growth and differentiation of the developing vertebrate retina. We show that the tissue develops by gradually increasing the average rate of differentiation and reducing the average cell cycle length. Moreover, this balance between differentiation and cell cycle duration increases the yield of production of terminally differentiated neurons, and it is required to achieve a well-defined exponential growth. We also show that a potential regulator of this balance is Hedgehog (Hh), promoting simultaneously cell cycle progression and cell cycle exit of RPCs. Our results represent a detailed and accurate quantitative characterization of retinal neurogenesis and how signals regulate the balance between proliferation and differentiation during the development of the vertebrate retina.

**Significance Statement:** The precise differentiation of neural progenitors during development ensures the correct cognitive, sensory, and motor functions of higher organisms. In this contribution, we characterize with quantitative precision the dynamics of growth and differentiation of the zebrafish developing retina during the first wave of neurogenic differentiation, using a combination of experimental/theoretical and computational tools. Our analysis reveals a progressive shift from proliferative to differentiative divisions, which correlates with cell cycle acceleration. Using small-molecule inhibition, we show that Hedgehog (Hh) signaling promotes both cell cycle exit and cell cycle progression, a balance required for well-defined exponential growth of the tissue.

## 1. Introduction

Cells sense, interpret, and respond to changes in their environment. During development, the precise interpretation of these external signals ensures the appropriate structure, dimensions, and composition of adult tissues and organs. Failures in this process are arguably one of the most frequently addressed topics in the field of Developmental Biology, since they can cause significant developmental disorders (1, 2), including physical and intellectual disabilities. In the context of neurogenesis in higher organisms, the balance between several of these morphogenetic signals sets the precise coordination between proliferation and differentiation of neural progenitors (3–7), ensuring the correct amounts of the different neuronal subtypes required for normal cognitive, sensory, and motor functions (8).

The developing retina is the most accessible portion of the vertebrate Central Nervous System, and an ideal organ for studying the dynamics and regulation of organogenesis in the context of neuronal differentiation (9, 10). The basic processes involved in the formation of the vertebrate retina and the specification of its various neuronal subtypes are well characterized (11). It starts with the evagination of the optic vesicle from the neural tissue in the early embryo. Next, the optic vesicle invaginates, forming a bilayered optic cup. The outer layer of the optic cup will become the retinal pigment epithelium (RPE), while the inner layer will generate the neural retina. Here, retinal progenitor cells (RPCs) undergo extensive proliferation and neurogenesis (12), sequentially generating six types of terminally differentiated neurons: retinal ganglion cells, horizontal cells, cone photoreceptors, amacrine cells, rod photoreceptors, and finally bipolar cells (13, 14). During this process, the vertebrate retina grows following a clear exponential dynamics in terms of cell numbers (15, 16), while cell density increases and the average cell and nuclear volume decrease (17).

Despite this conserved temporal organization and apparent simplicity, some key aspects of the formation and regulation of the developing vertebrate retina remain open: How do RPCs achieve this complex coordination between proliferation and differentiation in both space and time? How are these two processes coupled at the single-cell level? How do individual cells balance cell cycle exit and cell cycle progression to achieve this well-defined exponential growth dynamics at the tissue level?. How are these two processes balanced to achieve an optimal generation of differentiated neurons? Several of these important questions have been addressed previously, often based on qualitative changes observed in representative sections of the tissue. On the other hand, the complex organization of the vertebrate retina strongly complicates the definition of a single section that represents the processes in the entire tissue (18). Moreover, qualitative approaches may be sufficiently accurate when interpreting severe phenotypes, but in scenarios as complex as the vertebrate developing retina, they may potentially lead to inaccurate conclusions. This is why robust studies of the dynamics of neurogenesis in the vertebrate retina require quantitative data at the level of the whole tissue.

The quantitative characterization of biological processes at the level of single cells in highly dense three-dimensional tissues is a very challenging task. Many commonly used tools for automated 3D analysis work well when cells are well separated and defined, but struggle when applied to images of crowded tissues with little space between cells (17). In this direction, we recently developed a 3D image segmentation tool (OSCAR) specifically designed to quantify images of highly dense biological tissues acquired *in vivo* or *in toto* at suboptimal resolution (17).

Here, we adapt OSCAR to study with quantitative precision how cells in the developing vertebrate retina balance proliferation and differentiation to achieve the exponential growth of the tissue, and how key signal factors affect this balance. Our analysis reveals that the total number of cells during the first wave of neurogenesis in the vertebrate retina is defined by a reduction in average cell cycle length as the cells shift gradually from proliferative to differentiative divisions. Moreover, this combined modulation in both mode and rate of division ensures a well-defined exponential growth of the developing retina in terms of total number of cells. We also show that this dual modulation maximizes the number of neurons produced during this first wave of neurogenesis. We propose that Hh plays a major role in this balance by showing that it modulates the mode of division and cell cycle progression simultaneously. Overall, our study presents a quantitative characterization of the developing vertebrate retina and provides an accurate framework to study how system properties such as tissue growth and organization arise from processes at the level of single cells.

## 2. Results

### A. Exponential growth of the developing retina requires modulating the cell cycle length to compensate for terminal differentiation

In the context of population dynamics, the doubling of a population of cells at a given rate results in exponential growth dynamics. This type of growth is the expected solution in conditions of a pool of cells that proliferate at a constant rate (19). On the other hand, during the first wave of neurogenesis in the developing vertebrate retina, RPCs are also differentiating, meaning that some cells are continuously leaving the pool of proliferating cells. In these conditions, the typical doubling dynamics underneath every exponential growth may not be as trivial as initially expected.

To illustrate this and to study how this process of terminal differentiation may affect the growth dynamics of proliferating populations, we developed a simplified computational model. The model is based on agents that represent cells that perform tasks based on probabilities defined by the user (see Methods). In our model, agents can either proliferate (one cell undergoes mitosis and generates two identical cells) or differentiate (cells that exit the cell cycle and cease to proliferate). For simplicity, both processes are defined around mitotic events, labeling a cell division that produces two proliferative cells as a *pp* division, a cell division that produces two differentiated cells as a *dd* division, and a cell division that produces one proliferative and one differentiated cell as a *pd* division.

When we define these three options as probabilistic outcomes of a given division event, our agents in the model select between these three possibilities, so the probability to generate two progenitors is set by the value *pp*, the probability to generate two differentiated cells is set by the value *dd*, and the probability to generate one progenitor and one differentiated cell is set by the value *pd*. Since these parameters define probabilities, each one can take values from 0 to 1 (with *pp* + *pd* + *dd* = 1). Finally, if we only care about the outcome, and not about the value of each particular rate, we can define a new parameter *pp* − *dd* that takes values from 1 (all divisions are proliferative) to − 1 (all divisions are differentiative). The value *pp* − *dd* = 0 represents the homeostatic scenario where, on average, the pool of progenitors remains constant (Figure 1A). The other parameter of the model is the time between two mitotic events, represented as *T*.

**Fig. 1.**
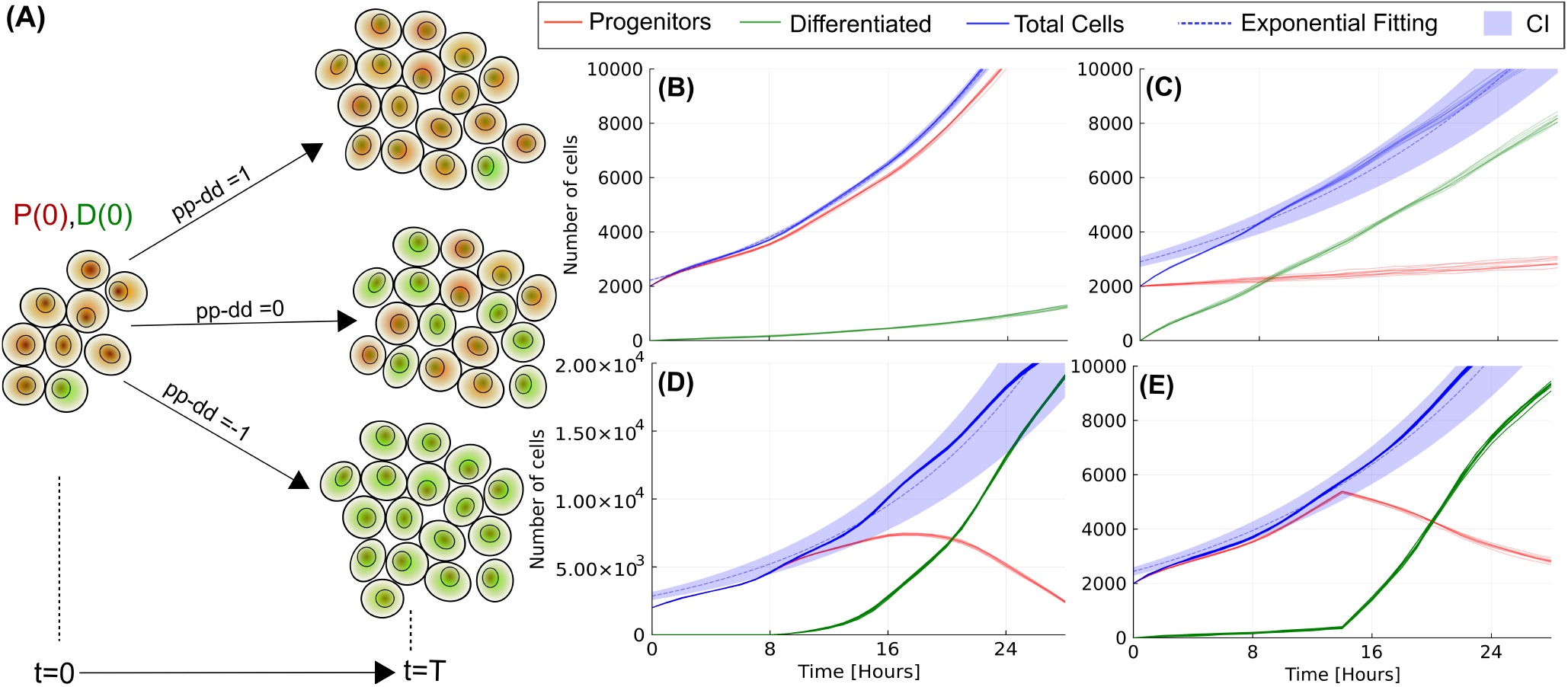
The dynamics of growth of a developing tissue depend on the balance between proliferation and differentiation. (A) Scheme of the numerical model of an initial population of progenitor cells (red) and terminally differentiated cells (green). After a cell cycle length, the progenitors divide to generate more progenitors or more differentiated cells based on the value of the parameter pp-dd (see Methods). (B) Dynamics of growth of the population (blue lines) for uniform *pp* − *dd* = 0.95 fits well to an exponential function (dashed blue line). (C) For values of the differentiation parameter uniform *pp* − *dd* close to zero (homeostasis in the progenitor population), the growth is mainly linear. (D) Dynamics when the differentiation parameter *pp-dd* changes gradually from 1 to −0.5 during the simulations. (E) Dynamics when the differentiation parameter pp-dd switches abruptly from *pp* − *dd* = 1 to *pp* − *dd* = −0.5 at *t* = 14 hpf. Ribbons in all panels of the figure represent the 95% confidence interval (CI) of the fitting. The centre for the error bands in all panels of this figure (dashed lines) corresponds to the average value of 10 independent numerical simulations. Parameters for the fitting are shown in Supplementary Table 3.

The starting point of the simulations is obtained from experimental data of the retina developing zebrafish at 20 hours post-fertilization (hpf) (17) in terms of cell numbers. The cell cycle length is fixed at around 6 hours, estimated also from experimental data (15, 16). The dynamics of growth in terms of the populations of progenitors (red) and differentiated (green) cells are plotted in Figures 1B-E for 28 hours, to mimic the duration of our previously published data (17). The model shows that when proliferative divisions dominate (*pp*− *dd* = 0.95, Figure 1B), the dynamics of growth of the full population (blue) fit well to an exponential (dashed blue line), as expected (goodness of fit is represented by the 95% confidence interval, in the form of blue ribbons around the fitted function). Figure 1C represents a situation of higher differentiation (*pp*− *dd ≈* 0, i.e., proliferation and differentiating divisions occur with similar probability). In these conditions, the dynamics do not follow the typical exponential dynamics (ribbons become larger), and the total number of cells fits better a linear growth.

On the other hand, it is well known that neurogenic tissues have an initial phase of expansion in the population of progenitors, followed by a phase of increased differentiation. This scenario is shown in Figure 1D, assuming a gradual monotonic shift between increased proliferation (*pp*− *dd* = 1) to increased differentiation (*pp*− *dd* = −0.5). In this case, the growth of the tissue does not fit an exponential function, as shown by the large confidence interval of the fitting. Finally, we test the effect of a sharp switch in the rates of *pp* and *dd* divisions, as we have shown that occurs during the differentiation of motorneurons of the vertebrate developing spinal cord (20), mediated by Sonic Hedgehog. Figure 1E has been obtained implementing a switch from *pp* − *dd* = 1 to *pp*− *dd* = −0.5 at t=14 hpf, showing again that the growth in number of cells does not fit an exponential function.

In conclusion, our simplified numerical model of a proliferating and differentiating cell population illustrates that, when differentiation becomes relevant (either via constant or variable rates), the exponential growth is not a trivial solution. Interestingly, these numerical predictions contrast with the well-defined exponential growth (17) observed for the developing retina from 20 hpf to 48 hpf (where terminal neurogenesis is known to be high). Therefore, our model suggests that exponential growth in terms of the total number of cells can only be achieved if other features of the process compensate for the effect of differentiation, such as the cell cycle length, the growth fraction, or the apoptosis rate.

### B. The identity of cells in the developing zebrafish retina can be established using differential protein expression

To study how the developing neural retina achieves its well-defined exponential growth despite undergoing differentiation, we designed a protocol to identify neural progenitors and terminally differentiated cells based on the Tg(atoh7:GFP) zebrafish line, combined with Sox2 immunostaining (see Methods). Atoh7 is a well-characterized member of the basic Helix-loop-Helix (bHLH) proneural transcription factor family expressed in terminally differentiating and differentiated cells (21, 22). Sox2 is a transcription factor that plays an essential role in the maintenance of both embryonic and adult neural progenitors (23).

Figure 2A shows the middle section of a zebrafish retina of the Tg(atoh7:GFP) line at 30 hpf stained for Sox2 (orange) and To-pro3 (grey, used for nuclear staining). Overall, cells expressing GFP have consistently reduced levels of Sox2 (examples marked with white arrows). Interestingly, some cells can be categorized as expressing intermediate levels of Sox2 and GFP (examples marked with yellow arrows), suggesting that both markers are not 100% exclusive. To further study this population of double-positive cells, we include immunostaining against Pcna, a crucial component of the replication and repair machinery commonly used as a marker of cellular proliferation (Figure 2B). Again, white arrows show that cells with high levels of GFP and low levels of Sox2 also show reduced levels of Pcna, suggesting that they are not actively proliferating (24). On the other hand, most cells with low levels of GFP and high levels of Sox2 also show overall high levels of Pcna (orange arrows), suggesting that they are actively proliferating. Interestingly, a subset of cells shows high levels of Sox2 and reduced levels of Pcna (illustrated by pink and blue arrows). Some cells in this subset (pink) correspond with cells undergoing mitosis (as illustrated by the TO-PRO-3 staining), in agreement with the reduction of Pcna in this phase of the cell cycle (25). The other subset (blue arrows) shows cells that are not differentiated but not actively cycling (26), suggesting that a percentage of progenitors may be in a quiescent state (not actively cycling). This subset of cells will become relevant later in the manuscript, when estimating the growth fraction of the progenitor population.

**Fig. 2.**
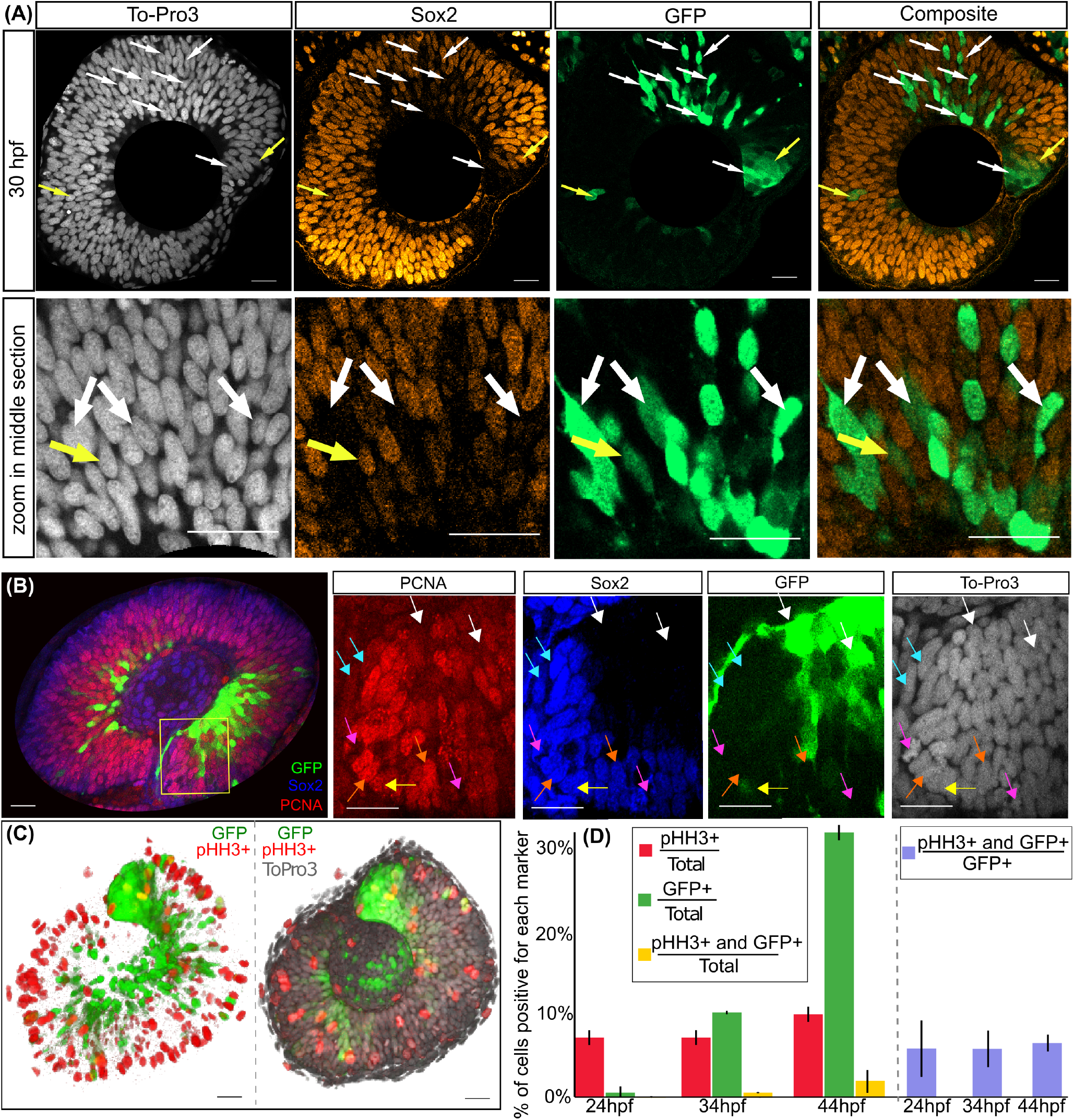
Characterization of the identity of the cells in the developing vertebrate retina based on differential protein expression. (A) Different channels of an image of a representative section of a developing retina at 30 hpf. A zoom of each image is shown below to illustrate that cells with high GFP expression correspond to cells that downregulate Sox2 (white arrows). A subset of cells (yellow cells) appears to express high levels of both markers. (B) Central slices of a zebrafish retina at 30 hpf co-stained with Pcna (red), Sox2 (Blue), and GFP (green). White arrows point to cells Pcna- and Sox2- and GFP+ (differentiated). Orange arrows point to cells Pcna+ and Sox2+ and GFP-(cycling progenitors). Blue arrows point to cells Pcna-, Sox2+, and GFP-(quiescent progenitors). Pink arrows point to mitotic cells negative for Pcna and GFP and positive for Sox2. Yellow arrows point to mitotic cells positive for GFP. (C) 3D-view of a zebrafish retina imaged in toto at 34 HPF stained with pHH3 (red), GFP+ (green), and To-Pro3 (grey). pHH3 staining appears in the whole retina (not only in the apical region) due to the 3D projection visualization. (D) Percentage of cells that are pHH3+ (red column), GFP+ (green column), and double positive for pHH3 and GFP (yellow column). In purple, the percentage of GFP+ cells that are both GFP+ and pHH3+ at different time points remains around 7%. Errobars correspond to the standard error of the mean value, from three independent repeats. Scale bars in all microscope images corresponds to 20 *µ*m.

Finally, we observed some cells that are positive for Pcna and Sox2 but also express GFP (marked by the yellow arrows), consistent with previous studies (11, 27, 28). To further confirm this, we perform immunostaining in the Tg(atoh7:GFP) line against Phosphohistone H3, a protein that is specifically phosphorylated during mitosis. The 3D reconstruction of an entire developing retina at 30 hpf is shown in Figure 2c (right panel contains the TO-PRO-3 channel also), confirming the existence of cells positive for both GFP and Phosphohistone H3 (in yellow). This is consistent with previous studies where authors monitored the activation of this Atoh7 promoter, reporting the existence of cycling GFP+ cells that divide to become photoreceptors, amacrines, retinal ganglion cells, horizontal cells, and even bipolar cells (24, 28). Next, we measure the percentage of cells positive for each marker (with an intensity value higher than an automated threshold) at three different developmental times. Results summarized in Figure 2D show that the percentage of mitotic cells remains stable around 8%, while the percentage of cells positive for GFP and double positive cells increases gradually. Interestingly, the percentage of double-positive cells in the population of GFP+ cells remains constant in the three time points tested (purple bars).

In conclusion, the combination of Sox2 and GFP allows us to define three different populations inside the developing vertebrate retina: cells with high levels of Sox2 and low GFP can be identified as progenitors (they are also Pcna+); cells with high levels of GPF and low Sox2 can be identified as terminally differentiated (they are also Pcna-); a small but non-negligible subset of cells with high levels of both GPF andSox2 that are also cycling (based on Pcna and PhosphoHistone3 immunostaining). Recent studies suggest that these cells emerge mainly from asymmetric divisions, resembling the intermediate progenitors identified in the mammalian neocortex (29).

### C. Automated quantification and characterization of full retinas at single-cell resolution using 3D segmentation and differential protein expression

In this section, we take advantage of the different levels of expression of Sox2 and Atoh7 to quantify the development of the neural retina based on the numbers of the different cellular subtypes. To do that, we will take advantage of our tool (17) for automated 3D segmentation (OSCAR) modified to classify cells based on two staining (see Methods).

In brief, OSCAR first segments and localizes all nuclei in the 3D image (based on nuclear staining) and then computes calculations based on the intensity levels of the other channels of the image. Typically, cells are classified as positive and negative for a given staining if their intensity is above and below a given threshold. In this case, the use of double staining allows us to classify cells without defining a threshold, resulting in more robust and accurate classification compared to single staining. This way, OSCAR computes the intensities of GFP (*I*_*GFP*_) and Sox2 (*I*_*Sox*2_) inside the volume occupied by each nucleus identified (illustrated in Figure 3A and explained in detail in the Methods Section) and classifies all cells detected as Progenitors (*I*_*Sox*2_ *>* 2 *× I*_*GFP*_), Differentiated (*I*_*GFP*_ *> ×* 2 *I*_*Sox*2_) or Intermediate Progenitors (1/2 < *I*_*Sox*2_*/I*_*GFP*_ *<* 2) (29). This two-fold intensity difference was chosen empirically as a good value for robust and accurate separation between the populations.

**Fig. 3.**
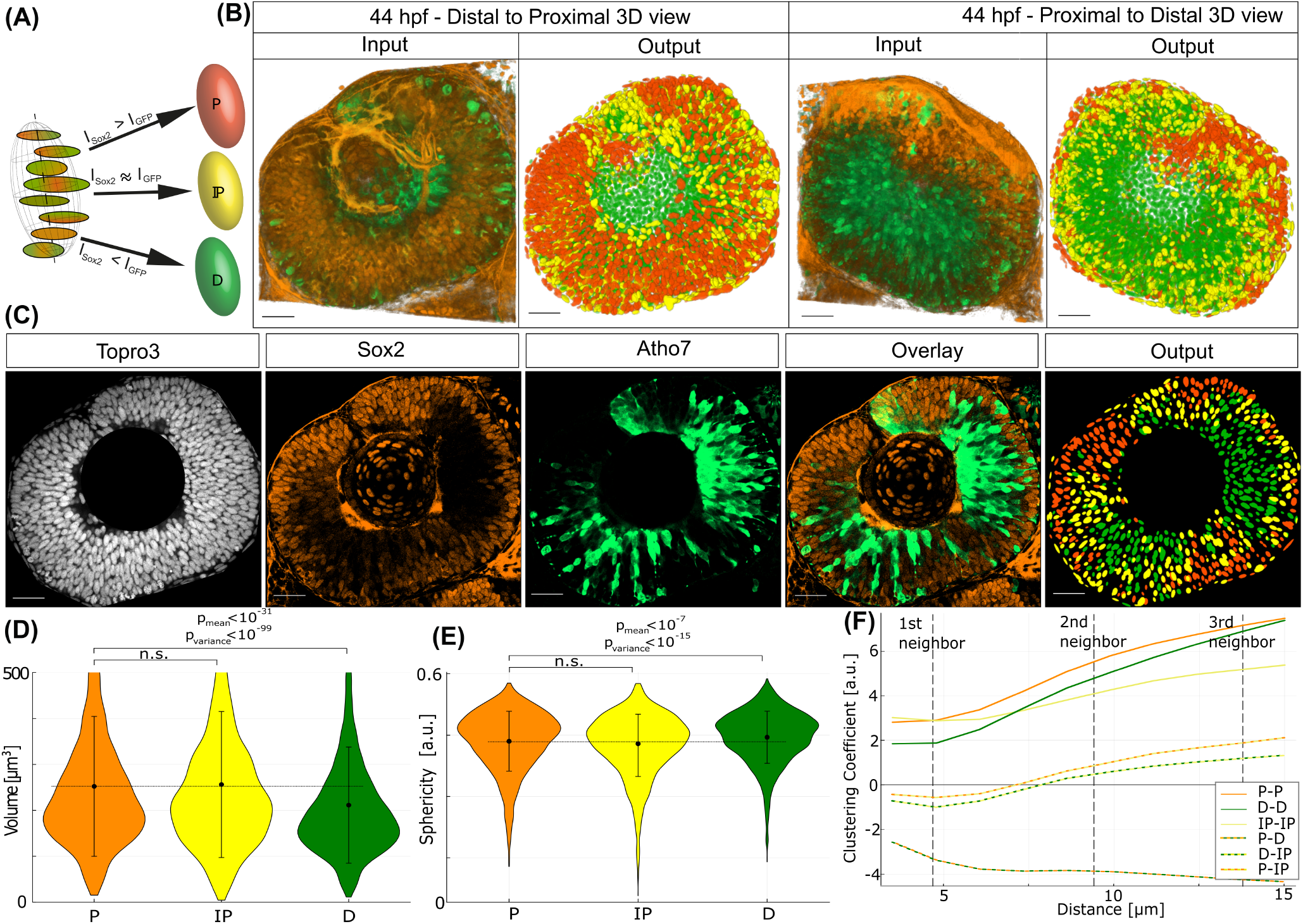
Segmentation and classification of single cells using OSCAR. (A) Scheme of the decision process to establish the identity of the objects detected, based on two complementary staining. (B) Example image of an entire zebrafish retina at 44 hpf viewed from outside (left) and inside (right) of the embryo (Sox2 and GFP channels). Output produced by OSCAR is shown to the right of each image, with each detected nucleus drawn as a 3D ellipsoid, color-coded based on its identity as P (orange), D (green), and IP (yellow). (C) Middle plane of the previous image, showing the different channels, the overlay of the Sox2 and GPF staining, and the corresponding plane in the output image (right) for comparison. (D) Quantification of differences in size between *P* (n=3500), *D* (n=1500), and *IP* (n=1500) for the same image. Statistical significance estimated using a Welch T-test (one-sided) for average values, F-test for variance (*p*_*mean*_ = 0.3 *×* 10^−31^, *p*_*variance*_ = 0.6 *×* 10^−99^). Error bars represent the 95% confidence interval (CI). (E) Quantification of differences in sphericity between *P* (n=3500), *D* (n=1500), and *IP* (n=1500) for the same image. Statistical significance estimated using a Welch T-test (one-sided) for average values, F-test for variance (*p*_*mean*_ = 0.2 *×* 10^−7^, *p*_*variance*_ = 0.9 *×* 10^−15^). Error bars represent the 95% confidence interval (CI). (F) Quantification of clustering (Besag’s L-function) of the different cell types. Scale bars in all microscope images corresponds to 20 *µ*m.

To illustrate visually the performance of OSCAR in this context, we show in Figure 3B an input image of a zebrafish retina at 44 hpf and its corresponding output, generated by drawing 3D ellipsoids with the shape and orientation detected for each nucleus, color-coded based on the identity of the corresponding cell (a movie showing all views of input and output is included as Supplementary Movie 1). Figure 3C shows the individual channels of the central confocal plane of the image, and the overlay of the Sox2 and GFP staining. The corresponding output of OSCAR of the same plane is also shown for comparison. A movie of all confocal planes for the input and output images is included as Supplementary Movie 2, where it can be clearly seen that the first planes are characterized by a majority of nuclei identified as *P*. As we enter the retina, *D* cells and *IP* become more abundant, and in the final planes deep inside the embryo, the majority of nuclei are identified as *D*.

This digital output of the 3D retina allows us to examine multiple features of the developing retina, such as morphological or clustering differences. Analysis of the nuclei size (Figure 3D) shows that cells identified as *P* and *IP* have larger nuclei than *D* cells, with statistical significance. This is consistent with the condensation of chromatin after the establishment of a terminal gene expression program (30).

In addition, the nuclei of the *D* cells are more homogeneous in size than the *P* and *IP* populations (illustrated by the variance as error bars), with high statistical significance. Focusing on the nuclei shape, the quantification shows a small difference between the three populations, estimated by the sphericity factor (Figure 3E). Again, the population of cells identified as *D* is more homogeneous in shape than the other populations, with high statistical significance (p-value of the variance is lower than 10^−15^ based on the F-test). Previous studies correlate nuclear shape with cell movement (31); therefore, a portion of the nuclei in the *P* and *IP* may elongate as a result of interkinetic nuclear migration (32). Finally, the output of OSCAR is used to estimate the clustering coefficient of the three populations using Besag’s L-function (see Methods), which returns positive values for clustering (objects of a particular type are closer together than expected when randomly distributed) and negative values for dispersion (objects of a particular type are further apart from each other than expected when randomly distributed). Results of this quantification (Figure 3F) show that the three cell types (solid lines) tend to clusterize at first and second closest neighbor distances. Interestingly, the clusterization coefficient of *IP* (yellow solid line) increases more slowly with distance, suggesting that *IP* clusters are smaller in size.

When considering the cross-clusterization (dashed double color lines), the analysis shows negative values for clusterization between *P* and *D* (green-orange line), evidencing that these two cell types form separate clusters in the 3D space. On the other hand, cross-clusterization of *IP* with both *P* and *D* (orange-yellow and green-yellow) is negative at short distances, but positive at around second neighbor distances, suggesting that small clusters of *IP* localize at the boundary between *P* and *D* clusters.

In conclusion, the combination of complementary Sox2/Atoh7 staining with our 3D automated segmentation tool allows us to process and quantify entire developing retinas at single-cell resolution with high accuracy. Our analysis shows that nuclei of the cells identified as terminally differentiated are smaller and more homogeneous in size and shape than the nuclei of actively proliferating cells (*P* and *IP*). Finally, the analysis of clusterization reveals a well-defined organization in the 3D space of clusters of *P* and *D* cells, which are clearly separated, and a clear tendency of *IP* to locate between these two clusters.

### D. The rate of differentiation of NPCs in the neural developing retina increases gradually concomitant with a decrease in cell cycle length

One of the main advantages of OSCAR is that it achieves high accuracy in segmenting objects in highly dense tissues much faster and with less computational requirements than Deep Learning-based Methods (17). Here, we use this advantage to measure the numbers of the different cell types in full retinas at many time points during the first wave of neurogenesis (33, 34).

The experimental approach is designed as follows (summarized in Figure 4A, and explained in detail in the Methods sections): (a) embryos of the Tg(atoh7:GFP) line are allowed to develop under standard conditions and are subsequently fixed at various developmental times, from 20 to 48 hpf; (b) they are then processed, cleared, stained, and mounted using in-house developed chambers; (c) retinas are imaged *in toto* using a confocal microscope; (d) images are then processed and analyzed using OSCAR to locate (nuclei channel) and establish identity (additional channels) of each cell based on differential protein expression; (e) output data is represented using 3D ellipsoids in the predicted location and color coded based on cell identity; (f) numbers of cells of each subtype are fitted using in-house algorithms to estimate the best fitting (Akaike Method) and the confidence intervals (Delta Method), explained in detail in the Material and Methods section.

**Fig. 4.**
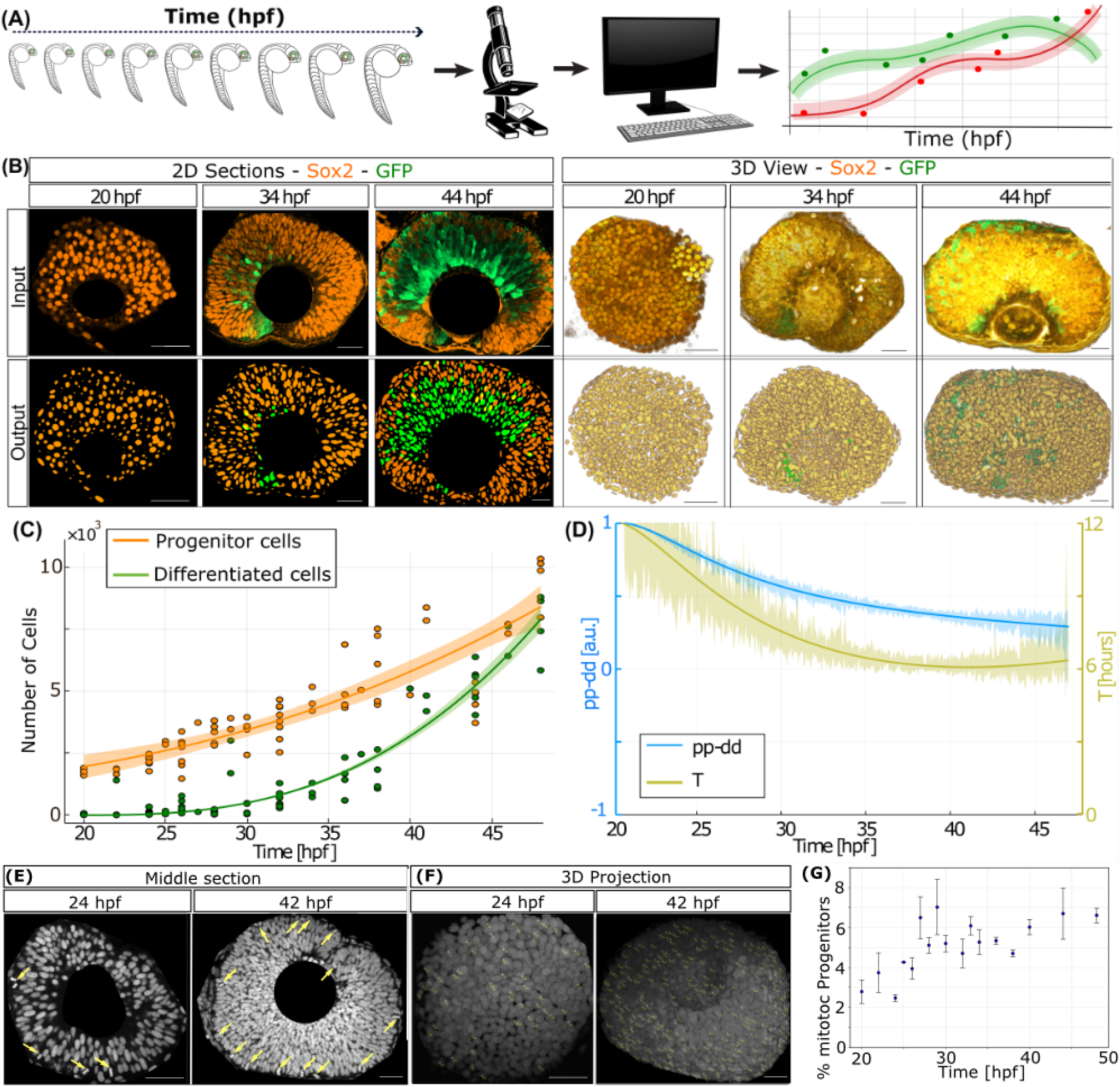
Quantification of the dynamics of proliferation and differentiation of the zebrafish retina. (A) Scheme of the experimental design (clipart from www.publicdomainvectors.com). (B) Representative sections and 3D-view of tissues stained with Sox2 (orange) and GFP (green) at different developmental stages. The corresponding digital representations are shown below. (C) Number of detected nuclei labeled as *P* (orange dots) and *D* (green dots) over time for 60 data points. The corresponding lines correspond to the fitting of the 60 individual retinas (obtained using the Akaike algorithm, see Methods). Solid lines correspond to the plot of the fitting. (D) Average *T* (dark yellow, right vertical axis) and *pp* − *dd* (light blue, left vertical axis) obtained from equations 1-2, when using as input the fitted curves *P* (*t*) and *D*(*t*). Ribbons in (C) and (D) illustrate the 95% confidence interval of the fitting, calculated using the Delta Method (see Methods). The centre for the error bands in C (solid lines) corresponds to the plot of the fitting (n=60, Akaike Method). The centre for the error bands in D (solid lines) corresponds to the value provided by the Delta Method when curves of panel C are used as input. (E) Middle section of two retinas stained with ToPro3 at 24 and 42 hpf. Yellow arrows mark mitotic events. (F) 3D projections showing all mitotic events in the entire retina. (G) Quantification of the percentage of progenitors undergoing mitosis in 57 entire retinas, showing an increase consistent with a decrease in cell cycle length. The centre of the error bands corresponds to the mean value of the values of each time point. Error bars represent the standard deviation. Scale bars in all microscope images correspond to 20 *µ*m.

To simplify the analysis, cells will be classified as proliferating (*P*) or terminally differentiated (*D*). These two subsets constitute the input of theoretical equations 1-2 (see Methods) to estimate the average mode and rate of division (intermediate progenitors are included as part of the proliferating population, since they are still actively cycling and dividing, see Discussion). Figure 4B illustrates examples of input images (first row) and the corresponding digital output (second row) of retinas at different hpf (left panels show the central plane of the retina, and right panels show the corresponding 3D view). Figure 4C plots the number of *P* (orange dots) and *D* (green dots) cells for around 60 full retinas. Optimal fitting for *P* cells (based on the Akaike algorithm using as input the number of progenitors for the 60+ retinas) is a sigmoidal function (orange line), consistent with a change in the growth rate of this population from high to low. On the other hand, the optimal fit obtained for the growth of the *D* population (based on the Akaike algorithm using as input the number of differentiated cells for the 60+ retinas) is a power law (green line), consistent with the accumulation of these cells as they are continuously produced. Percentages of *P, D*, and *IP* cells are plotted as Supplementary Figure 1, showing that approximately 100% of cells are identified as *P* (orange) at initial time points. As time progresses, the percentage of cells identified as *D* (green) and *IP* increases (yellow, representing around 15% of cells at 48 hpf).

Next, the fitted values for *P* and *D* are used as input for a set of analytical equations based on a branching process formalism developed by our group (35) (See Methods Section for details). In essence, equations 1-2 provide values for the difference between rates of proliferative and differentiative symmetric divisions (*pp* − *dd*), as well as the average cell cycle length (*T*) of cycling cells, with temporal resolution and higher accuracy than BrdU or EdU-based methods (7).

Parameter *γ* in the equations represents the growth fraction (i.e, the rate of *P* cells that are actively cycling), which takes values from 1 (all progenitors are actively cycling) to zero (all progenitors are quiescent). The value of *γ* is estimated here by calculating the ratio of Sox2+ that are also Pcna+ at three time points. Our data suggests that the growth fraction starts at *γ* = 1.0 at 20 HPF, consistent with previous studies based on BrdU incorporation (36). Next, it decreases slightly (*γ* = 0.85) at 35 HPF, before rising again to *γ* = 0.95 at 48 HPF (estimated based on 3D quantification of three independent replicas per time point). Based on this, we estimate that *γ* changes smoothly over time following these values, using spline interpolation.

The final parameter required is *∅*, which represents the rate of apoptosis in the population of progenitors. To estimate this, we use a dual approach based on quantification of immunofluorescence against the active form of Caspase 3 and manual identification of piknotic nuclei (both combined with Sox2 immunostaining) Estimation of the number of cell undergoing apoptosis at t=20, t=35, and t=48 HPF (three independent replicates per time point) shows a similar value of around 0.1% of the progenitors can be identified as undergoing cell death, in agreement with previous studies (16, 37, 38) (see Supplementary Figure 2A as an example of a cell showing Active Caspase 3 immunostaining and piknotic nuclei). To translate this value as a rate of apoptosis (number of cells that undergo apoptosis per time point), we estimated the average time interval for piknotic nuclei to be detected in time-lapse movies as 3.5 hours. This results in an average rate of apoptosis in wild type embryos of *∅* = 0.00045Ė stimation of the effect of this rate of apoptosis in the final population of progenitors and differentiated cells in the retina is illustrated in Supplementary Figure 2B using numerical simulations.

Using these values for*∅* and *γ*, along with the fitted values for *P* and *D* from 20 to 48 hpf, the average mode of division *pp* −*dd* (Figure 4D blue line, left vertical axis) predicted by equation 1 suggests a shift from mainly symmetric proliferative divisions (*pp* − *dd ≈* 1 at 20 HPF) to increased differentiation of RPCs (*pp* − *dd ≈* 0.35 at 48 HPF). Concurrently, equation 2 predicts that the average cell cycle *T* (Figure 4D yellow line, right vertical axis) accelerates monotonically from an initial value of *T ≈* 12 hours at 20 HPF to *T≈* 6 hours at 48 HPF. To validate this acceleration in cell cycle length, we quantified the percentage of progenitors in M-Phase in 30 retinas at different time points (assuming that the length of M-phase is constant, a decrease in the other phases of the cell cycle should result in an increase in the percentage of progenitors in M-phase). Representative examples of sections and 3D projections of retinas at two time points are shown as Figure 4E, where yellow arrows are drawn to illustrate mitotic events (4 at 24 hpf, 14 at 42 hpf). The 3D projection of the entire nuclei Figure 4F, shows all mitotic events found in all sections of the image. Data of 30 full retinas at different time points shows an increase in the percentage of progenitors in M-phase as development proceeds, with values at initial time points around 3% and at later time points around 6%. This increase in the percentage of progenitors in mitosis is consistent with the acceleration form 12 hours to around 6 hours in cell cycle length, measured using the branching equations.

In conclusion, the combination of experimental data *in toto* with automated 3D segmentation allows us to assess the balance between proliferation and terminal differentiation during the initial wave of neurogenesis in the zebrafish retina, providing also the measurement of key morphological features at the tissue level as well as at the single cell level. The values obtained for progenitors and differentiated cells, when used as input for the branching equations, show that both the average mode of division and the average cell cycle length change simultaneously during the first wave of neurogenesis of the vertebrate retina.

### E. Cell cycle acceleration improves the exponential growth and increases neuronal production

The previous experimental approach allows us to obtain quantitative measurements of the average mode and rate of division using data from entire developing retinas. By construction, these values are also the input values required in the agent-based model developed in the previous sections. This way, if correct, numerical simulations of the model using as input the values of *pp* − *dd* and *T* predicted by the branching should reproduce the experimental data. This is shown in Figure 5A (using the same value of *γ* and *∅* as used for the branching equations), where the output of 10 numerical simulations (solid green and red lines) is plotted on top of the experimental data (dashed green and red lines). Based on the good agreement between model and experiment, we can conclude that the values of *pp* − *dd* and *T* measured correctly reproduce the overall growth and differentiation dynamics of the tissue.

**Fig. 5.**
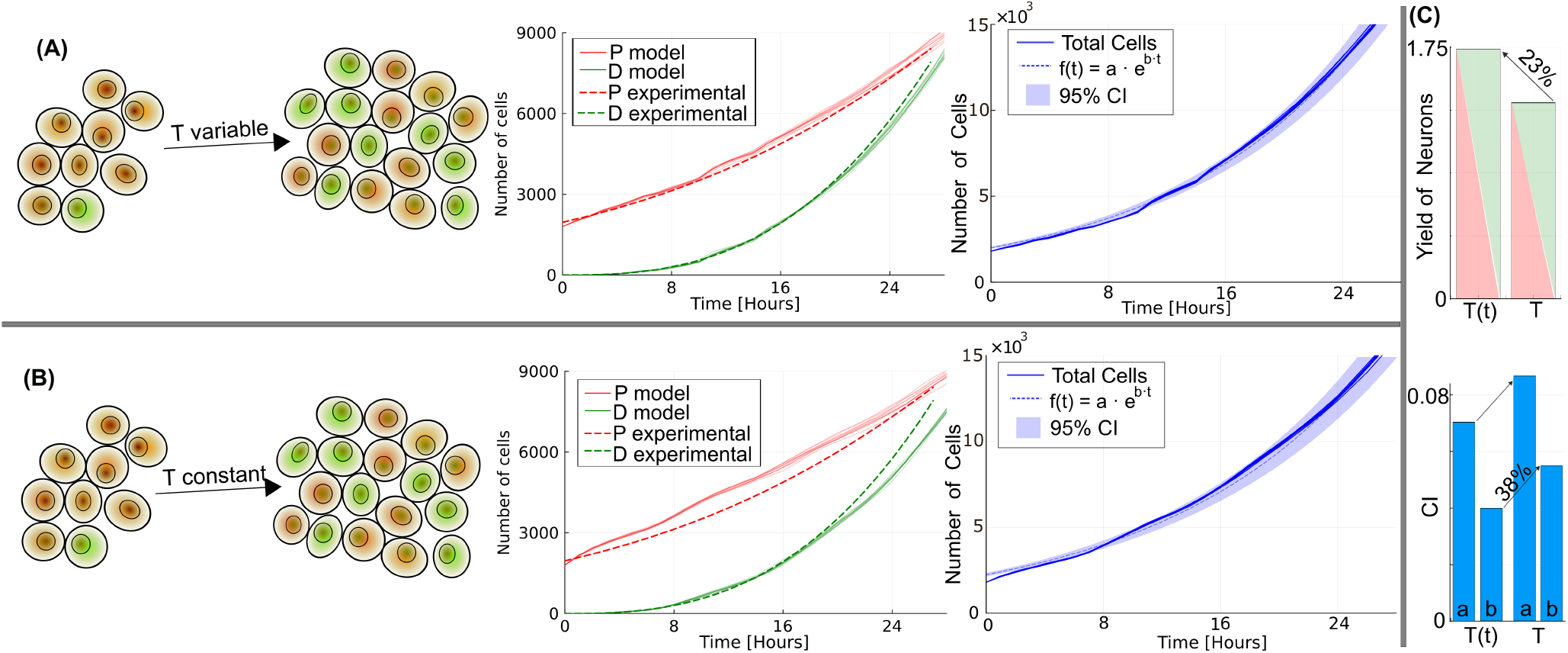
Cell cycle acceleration increases the yield of neurons and compensates for terminal differentiation to achieve exponential growth. (A) Numerical simulations of the model using as input the mode of division *pp* − *dd*, the cell cycle length *T*, the growth fraction *γ*, and the apoptosis rate *ϕ* measured in the experiments. The simulations (solid lines) reproduce well the values of the experiment (dashed lines). When computing the total number of cells (solid blue lines on the right panel) of 10 independent simulations fits well to an exponential curve (ribbons illustrate the 95% confidence interval, calculated using the nonlinear least-squares method. The centre of the errorbands (dashed lines) corresponds to the average of 10 independent simulations). (B) Numerical simulations of the model were performed using as input the mode of division measured in the experiments, and a constant cell cycle length of 8 hours. The number of cells predicted by the model (solid lines) does not follow the experimental data (dashed lines). The total number of cells (solid blue lines on the right panel) presents a worse fit to an exponential curve, compared to the plot above (variable T). (C) Decrease in neuronal yield (neurons produced by each progenitor on average) and increase in the 95%CI in the two parameters of the exponential fit. Dashed lines corresponds to the average of 10 independent simulations. Parameters for the fitting are shown in Supplementary Table 3. The centre of the errorbands (dashed lines) in all panels of this figure corresponds to the average of 10 independent simulations).

Next, we can use the model to test the effect of the variations in the cell cycle length of the progenitors observed in the experiments. To do that, we perform simulations using the same values of *pp* − *dd* and *γ*(*t*), but using a constant value of *T* =7 hours (to obtain the same number of total cells at the end of the experiment). The dynamics in this scenario are plotted in Figure 5B, showing that the system generates the correct number of *D* but a higher number of *P* at intermediate time points (around 16 hours). On the other hand, the model achieves the correct final values of *P*, but results in a smaller number of *D*. This suggests that the strategy of variable cell cycle (Figure 5A) is capable of generating a higher number of differentiated cells from a smaller number of progenitors, compared to a situation of fixed cell cycle length. Quantification of the yield (Figure 5C, in terms of the number of *D* cells at the final time point (t=28 h) per progenitor cells at intermediate time points (t=14 h)) shows that each progenitor results in 23% more neurons (from 1.42 to 1.75 neurons per progenitor on average) in conditions of variable *T*.

In terms of the total number of cells (blue lines), no significant differences are observed between both situations (variable *T* in Figure 5A and fixed *T* in Figure 5B). On the other hand, when these values are fitted to exponential curves (dashed lines), the system with variable *T* (Figure 5A) results in a much better fit, represented here by the width of the ribbons, which illustrate the 95% confidence interval of the fitting. This is clearer if we plot the value of the 95% CI of both parameters of the exponential (Figure 5C), with a 38% increase in the growth rate of the exponential. The value of the square root of the square of the residuals also shows a 31% increase (from 1015 for variable *T* to 1326 for fixed *T*).

In conclusion, the model validates that the experimental approach based on OSCAR and the branching equations provides the correct values for the average mode and rate of division. In addition, comparing with a situation of a fixed cell cycle, the variable cell cycle increases the yield of differentiated cells from fewer progenitors. Moreover, this modulation in the length of the cell cycle results in well-defined exponential growth in terms of the number of total cells, as observed experimentally.

### F. Hh signaling promotes cell cycle exit and cell cycle progression during the first wave of differentiation in the developing vertebrate retina

A potential explanation for the dual modulation of mode of division and cell cycle length is the regulatory processes reported to affect both differentiation and cell cycle dynamics. A good candidate is Hedgehog (Hh) signaling, one of the most well-characterized regulators of developmental processes (4–6, 20), involved in the genesis of most organs in higher organisms. One of the most extensively studied roles of Hh is as a major driver of patterning and differentiation of neural progenitors during embryogenesis (20, 39–43). In addition, Hedgehog has been linked to the promotion of cell proliferation by inducing the transcription of Cyclin D and Cyclin E via the Gli transcription factors (44).

In the developing vertebrate retina, Hh is involved in a positive feedback loop that generates a traveling front (45) of Hh expression that starts at the ventro-nasal region and spreads centrifugally to organize the neurogenesis across the epithelial sheet (9, 46–49), similarly to the propagating wave of differentiation across the eye primordium in Drosophila (4–6, 50).

To characterize the specific role of Hh during retinal development, zebrafish embryos from the Tg(Atoh7:GFP) line are cultured in the presence of Cyclopamine (51), a small molecule inhibitor that selectively binds to the Hh receptor Patched and interrupts Hh pathway activity (see Methods). To estimate the efficiency of inhibition, we quantify images of *in situ* hybridization for Patch1 (the receptor of Hh that is also a target of the pathway). Figure 6B-C shows around a 10X decrease in Patch1 mRNA levels (FGF8 is used as a control for the quantification).

**Fig. 6.**
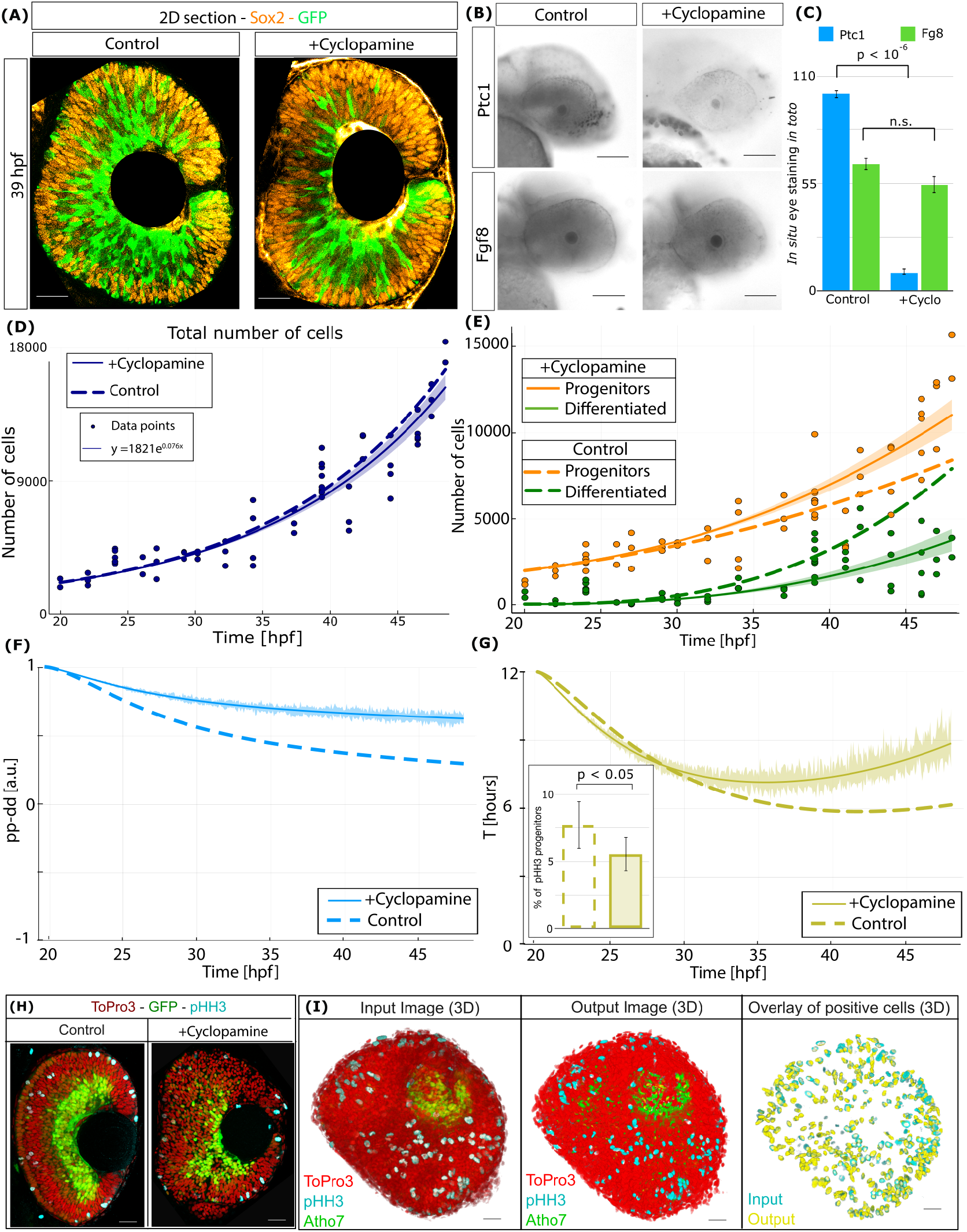
Hh promotes differentiation and cell cycle progression in the developing vertebrate retina. (A) Representative sections of tissues stained with Sox2 (orange) and GFP (green) at 39 HPF for control and +cyclopamine treated embryos. (B) *In situ* hybridization for Patch1 at 37 HPF for control and +cyclopamine conditions. *In situ* for FGF8 is used as a quantification control. (C) Intensity of *in situ* staining measured *in toto* for zebrafish retinas showed statistically significant differences in Patch1 (Student’s t-test, one-sided, *p* = 0.2 *×* 10^−6^), while non-significant for FGF8. (D) Total number of cells for 50 embryos treated with cyclopamine and fixed at different HPF. The solid line corresponds to the exponential fitting. Ribbons correspond to a 95% confidence of fitting. Fitting for control conditions (dashed line) is added for comparison. (E) Number of cells identified as progenitors (orange dots) and differentiated (green dots) over time in +cyclopamine-treated embryos. Solid lines correspond to the fitting for progenitors and differentiated cells (obtained using the Akaike method, explained in the Methods Section. Fitting for control conditions is added (dashed lines) for comparison. (F-G) Average *pp* − *dd* (panel F, blue) and *T* (panel G, yellow) obtained from equations 1-2, when using as input the fitted curves *P* (*t*) and *D*(*t*). Ribbons illustrate the 95% confidence interval of the fitting, calculated using the Delta approximation (see Methods). The insert shows the percentage of pHH3+ progenitors for control and cyclopamine-treated embryos at 45 HPF (statistical significance estimated using the Welch’s t-test with a one-tailed directional hypothesis, *p*=0.03). (H) Central confocal section of zebrafish retina at 45 HPF showing TO-PRO-3 (red), GFP (green), and pHH3 (blue) immunostaining for control and +cyclopamine conditions. (I) 3D-view of a representative input image (left panel), the corresponding objects colored by their identity, and an overlay of only objects marked as positive for pHH3 (yellow) on top of the immunostaining for pHH3+ (blue). Erro-bars correspond to the standard error of the mean value, from three independent repeats. Scale bars in panels A,H,I corresponds to 20 *µ*m. Scale bars in panels B corresponds to 100 *µ*m.

Figure 6A displays the middle plane of a zebrafish retina at 39 HPF from both a control embryo (left) and a +cyclopamine treated embryo (right), stained with Sox2 (orange) and anti-GFP (green). Comparison of these sections suggests that cyclopamine-treated retinas exhibit a similar size but a smaller proportion of GFP+ cells. However, this difference could also stem from variability between embryos or sections of the same retina, as the proportions of progenitors and differentiated cells are highly dependent on the plane selected (18).

To determine the statistical significance of the differences induced by Hh inhibition, we quantified approximately 50 embryos at different HPF (Figure 6D, solid line), revealing a trend that aligns well with an exponential curve, with a negligible reduction in size compared to the control conditions (dashed line). Conversely, we observe a statistically significant increase in the numbers of progenitors (Figure 6E, orange dots, orange solid line) and a reduction in differentiated cells (Figure 6E, green dots, green solid line) compared to the control (dashed lines, reproduced here to facilitate comparison), particularly at later time points.

These numbers are then utilized as input for equations 1-2 (cell death and quiescence remain equivalent between control and +cyclopamine conditions), and the average mode and rate of division (solid lines) are compared to control conditions (dashed lines). The output for the mode of division (Figure 6F, blue) shows a gradual shift towards more proliferative divisions, compared to the control.

Interestingly, this increase in the rate of proliferative divisions is expected to generate more progenitors, leading to larger retinas (as more cycling cells would produce more progeny). This discrepancy is clarified when we focus on the cell cycle length of progenitors (Figure 6G, yellow solid line), which is now, on average, slower than in control conditions (yellow dashed line), particularly from 35 hpf onwards (based on non-overlapping 95% confidence intervals, yellow ribbons, obtained using the Delta Method). This longer cell cycle balances the increase in progenitors after Hh inhibition, resulting in a similar exponential growth as in the control conditions.

To further validate this shift in cell cycle length, we quantified the percentage of cells identified as progenitors (GFP-) that are also labeled as positive for pHH3+ for control and +cyclopamine conditions at 45 HPF (assuming that the length of M phase is constant, an increase in the other phases of the cell cycle should result in a decrease in the percentage of progenitors in M-phase). Figure 6H shows the mid-section of retinas for the two conditions, showing a clear reduction in differentiation (green) and apparent reduction in pHH3+ staining. To test if this change is statistically significant and representative of the whole tissue, we processed and quantified using OSCAR around 10+ retinas. Figure 6I shows an example of a 3D view of input, output, and overlay of pHH3+ objects (yellow) over the input image (blue), showing very good agreement between the output and input. Quantification of these images (insert in Figure 6G) shows a reduction in +cyclopamine (solid line) versus control (dashed line) in the percentage of pHH3+ progenitors that is statistically significant. To further validate the values of *pp-dd* and *T* in these conditions, we run numerical simulations of the agent-based model using them as input to the model. The good fit between the model prediction and the experimental data (Supplementary Figure 3) suggests that the values extracted for the mode and rate of division are correct.

In conclusion, Cyclopamine treatment in developing zebrafish retinas diminishes differentiation by promoting more proliferative divisions. Simultaneously, the cell cycle length increases to compensate for the higher number of progenitors, resulting again in exponential growth. This effect is consistent with a role of Hh in regulating neurogenesis by promoting the cell cycle exit, but also cell cycle progression.

## 3. Discussion

The study of biological processes with quantitative precision provides advantages that go beyond the simple characterization of a system using numbers. For instance, this level of accuracy allows us to compare changes between different experimental conditions, and to unveil mild but relevant differences between phenotypes that may not be noticed by eye inspection. In the context of drug treatment, it provides us with the possibility to characterize effects at low doses (as we do here with cyclopamine), reducing potential off-target interactions.

Some of the quantitative data reported here aligns well with previous studies, providing a solid validation that may reduce the controversy between studies that suggest contradictory results. For instance, on the topic of the link between cell cycle length and division outcome, previous works reported lengthening of the cycle for differentiated divisions (52–62), shortening (20, 36, 63–67), or even no dependence (35, 68, 69). Focusing specifically on the developing retina, a detailed revision of the topic of interplay between mode and rate of division in mouse and rat (62, 70–73) seems to follow different dependencies than in zebrafish and xenopus (36, 67, 74–77). Our analysis shows a clear decrease in the cell cycle length as differentiation increases, both in control conditions (*T* shortens as *pp* − *dd* becomes smaller) and after cyclopamine treatment (*T* is longer and *pp* − *dd* increases), in agreement with the literature in the zebrafish and Xenopus model (67). In addition, our measurement of the average length of the cell cycle (Figure 4D), taking the full progenitor population, agrees with values measured using single-cell tracking from 4 to 11 hours (78). In addition, our data is in agreement with previous work by the lab of Stephen Easter (36), showing a clear cell cycle acceleration taking place from 20 hpf to 29 hpf, reaching average values around 8 hours (also consistent with our data). Our analysis shows that this acceleration continues as differentiation increases to reach a value of around 6 hours, and is maintained at least until 48 hpf.

Interestingly, the dual modulation of cell cycle and mode of division increases the number of neurons being generated per progenitor, compared to a situation where the cell cycle length does not accelerate as neurogenic divisions increase. This may potentially play a role as an evolutionary strategy to maximize the yield of neurons from a small pool of progenitors, in the same way as fast cycling Transit-Amplifying Cells (TACs) in the mammalian (79) and invertebrate (80) brain.

A very recent study (81) shows that mice mutants with disrupted Hh signaling show retinas of similar size to control embryos (in agreement with our study). In addition, they show a significant reduction of cells in mitosis in Smo mutants, similar to our data. On the other hand, they propose a model where, in the absence of Smo, RPCs exit the cell cycle prematurely, which is again not consistent with our data in zebrafish, but consistent with previous studies in mouse models.

In terms of the balance between proliferation and differentiation, recent studies have focused on the dynamics of differentiation waves (24, 82) and the subsequent translocation of cells post-differentiation (83). Based on these and other studies, differentiation appears as the most relevant process at later time points, based on the eventual dominance of differentiated cells over progenitors. Interestingly, our results indicate that despite this, division events always produce more progenitors than differentiated cells (*pp* − *dd >* 0, progenitors are always produced at a higher rate than differentiated neurons) during this first wave of neurogenesis. This apparent discrepancy arises since progenitors continue to cycle, while differentiated neurons accumulate over time, eventually becoming the most abundant type.

Many previous studies link Hh to cell cycle exit, and others to cell cycle progression. Our analysis based on the branching formalisms allows us to measure both independently, showing that Hh in the developing retina plays a dual role in promoting both cell cycle exit and cell cycle progression. This scenario aligns with the role of Hh in transitioning cells from a slow-cycling progenitor to a faster-cycling progenitor more ready to terminally differentiate (84). This dual role explains the minimal changes in the overall size of the retina observed in our experiments. Typically, a simple reduction in differentiation following Hh inhibition would lead to an increase in cycling progenitors, potentially resulting in larger final retinas (an outcome inconsistent with previous observations (85) and our own data). However, combining this effect with an increase in cell cycle length compensates for the higher number of progenitors, resulting in retinas of similar size.

Previous studies propose that Atoh7+ progenitors in the zebrafish retina emerge mainly from asymmetric divisions of Atoh7-progenitors, and that they cycle just once more to generate two differentiated cells (29). Based on this scenario with a different mode of division for Atoh7+ and Atoh7-progenitors, our analysis could benefit from treating these two populations as separate. Unfortunately, solving a branching process formalism analytically with two types of progenitors (*P* and *IP*) is not possible (86). As a consequence, only the average *T* and *pp* − *dd* for all progenitors can be extracted from the changes in cell numbers. In addition, due to the mathematical equivalence between *pd* and *pp* plus *dd* division (35), the branching theory cannot be used to distinguish between asymmetric and symmetric divisions (see Supplementary Figure 4).

Interestingly, previous studies show that these progenitors positive for Atoh7-GFP have a longer cell cycle (87) than Atoh7-GFP negative progenitors (16). This contrasts with our findings that show that cell cycle accelerates concomitant with an increase in the amount of these Atoh7-GFP progenitors (Supplementary Figure 1). A potential explanation of this discrepancy is that the amount of Atoh7-GFP precursors is small compared to the rest of cycling progenitors, as shown by our analysis.

Another important limitation of these types of analysis based on population averages is that they may mask very interesting data regarding cell-to-cell variability as well as single-cell heterogeneity. Several previous studies have focused on clonal analysis or time-lapse trajectories of single cells, providing relevant information on these topics (12, 14, 28, 88– 90). Here, we approach the development of the retina with a *systems* perspective, based on our previous contribution that showed that the entire dynamics of development of a proliferating and differentiating tissue can be captured by the average mode of division, the average rate of division, and the growth fraction (7, 35). Therefore, our approach is designed to capture the emergent behavior at the tissue-level and not the specific single-cell behaviors that produce this average.

It would be interesting to measure if the acceleration of the cell cycle length is maintained after 48 hpf, or if it slows down as in the mammalian retina. Unfortunately, our approach is limited to time points before 48 hpf due to two reasons: (a) quantification in 3D of entire retinas becomes unreliable after this time point, due to sample thickness and light scattering, (b) a subset of Horizontal cells arising after 48 hpf show positive staining for both Atho7 and Sox2 (so our system will erroneously identify them as cycling).

A main drawback of small-molecule treatment is non-specific interactions, specially when used in high concentrations (47, 91). In this direction, our quantitative approach allows us to study the cyclopamine effect at low doses, where binding specificity towards its targets is supposed to be optimal. On the other hand, a main advantage compared to genetic approaches is that it allows us to apply inhibition at a very specific time point, constituting a good strategy to study the direct effect of a signaling molecule after a particular developmental stage. In the case of Hh, this is specially relevant, due to its role in eye vesicle patterning, specification, and morphogenesis, (92, 93), processes that take place hours before neurogenesis.

## 4. Conclusions

In this contribution, we present a novel experimental/computational/theoretical framework specifically designed to characterize the developmental dynamics of the vertebrate retina with unprecedented accuracy. The framework enables us to examine various key aspects of the initial wave of differentiation, including growth, composition, organization, and dynamics of proliferation and differentiation. We show that the cell cycle length shortens as differentiation increases, and that this dual modulation results in a higher yield of neurons and a more defined exponential growth in terms of total cell numbers. We propose that the vertebrate retina develops by following a strategy that optimizes for the rapid production of neurons while simultaneously maintaining a stable, consistent, and predictable growth dynamics. This interplay between mode and rate of division is mediated by Hedgehog signaling, actin as a promoter of cell cycle exit, and cell cycle progression.

## Materials and Methods

### A. Animals and Experimental procedure

All experiments were conducted in accordance with relevant institutional, national, and/or international guidelines and regulations for animal research. (permit number PROEX 347.4/23, issued by the Dirección General de Agricultura, Ganadería y Alimentación CONSEJERíA DE MEDIO AMBIENTE, AGRICULTURA E INTERIOR de la Comunidad de Madrid, Spain).

Experiments are conducted using embryos from the Tg(atoh7:GFP) zebrafish line, which has been genetically engineered to express GFP as a marker for Atoh7 expression (28). The animals are maintained and bred according to established protocols (94): (a) Adult zebrafish are maintained at 28°C in specialized couvettes in the fish facility with all permits and constant supervision. To obtain fertilized eggs, adults are placed in mating couvettes overnight, with males and females separated. (b) The barrier is removed the following morning, and the presence of eggs is checked every 20 minutes to accurately determine the date of birth based on visual inspection (95). (c) Fertilized eggs are collected and cultured in fresh E3 medium 1X (5 mM NaCl, 0.17 mM KCl, 0.33 mM CaCl2, 0.33 mM MgSO4, Methylene Blue), supplemented with phenylthiourea (PTU) at 0.003% from 20 hours post-fertilization (HPF) to prevent pigmentation, with media replacement every 24 hours. (d) Sets of 20 embryos at similar HPF are transferred to p100 Petri dishes and placed in an incubator at 28°C. (e) Pronase (Merck, CAS-No 9036-06-0) at a concentration of 1:100 in E3 1x medium is added at 20 HPF to remove the chorion (96). (f) Subsequently, embryos are divided into sets of 5 in 6-well plates, and either DMSO (control) or cyclopamine at 0.1 mM diluted in DMSO (Merck, 239806) is added. (g) Embryos are collected at developmental stages between 24 and 48 HPF and fixed overnight at 4°C in a 10% Formalin solution (Sigma, HT501128). (h) Subsequently, embryos undergo dehydration using a series of methanol dilutions (0%, 25%, 50%, 75%, and 100%) and then stored at −20°C for a minimum of overnight and a maximum of six months, following a protocol adapted from ref. (97).

### *B. in toto* immunostaining

(a) Re-hydration of embryos is conducted using the previous serial dilutions of methanol, but in reverse order. (b) All subsequent solutions are prepared in PBS 1X with 0.6% Triton to enhance the permeability of antibodies and dyes. (c) Embryos undergo incubation in proteinase K (10 ug/ml) for a specified duration (refer to Supplementary Table 2) to facilitate penetration of antibodies and dyes. (d) The reaction is halted by brief exposure to 10% Formalin (Sigma, HT501128) for 20 minutes at room temperature. (e) Embryos are washed three times with PBS 1X and 0.6% Triton. (f) Subsequently, embryos are incubated in a blocking solution (10% FBS in PBS 1X and 0.6% triton) overnight at 4°C to prevent non-specific antibody binding. (g) Embryos are incubated with the corresponding primary antibodies, diluted in a solution of 2% FBS in PBT 0.6%, overnight at 4°C. (h) The primary antibody solution is washed three times for five minutes each. (i) Embryos are then incubated in a solution of 2% FBS in PBT 0.6% with secondary antibodies and nuclei staining for 1 hour at room temperature. (j) Finally, embryos undergo three washes with PBT 0.6% followed by PBS.

The primary antibodies used are: chicken anti-GFP (1:1000, Abcam ab137827), rabbit anti-Sox2 (1:1000, GeneTex GTX124477), mouse anti-Pcna (1:100; Invitrogen, MA5-11358), and rabbit anti-PH3 (1:250; Merck-Millipore, 06-570). The secondary antibodies used are: goat anti-Chicken-488 (1:500; ThermoFisher, A-11039), goat anti-mouse-55 (1:500; ThermoFisher, A-21157) and donkey anti-rabbit-555 (1:500; ThermoFisher A-31573). DNA is stained with the intercalating nuclear agent 647 nm fluorophore To-Pro3 (1:500; ThermoFisher, T-3605) for all samples.

As a positive control for Active Caspase 3 immunostaining, we present in Supplementary Figure 2 a zebrafish retina at 52 HPF (when apoptosis becomes non-negligible) stained against Active Caspase 3 (37).

### C. In situ hybridization *in toto* protocol

To quantify the efficacy of Hh pathway inhibition by Cyclopamine, the integrated density of the Patch1 staining signal was measured across the entire 3D volume of the retina for each embryo. This value was then normalized to the signal from the FGF8 control probe in the same embryo to account for variability in staining and imaging. The protocol for the *in situ* hybridization was performed as follows:

#### Day 1: *Hybridization*

After fixation (Formalin 10%), dehydration and rehydration in MeOH, whole embryos are washed with PBS+Tween10%(PTW) 4 times for 5 minutes. Permeabilization is performed with Proteinase K at 10 *µ*g /mL (incubation time similar to the immunofluorescence assay, Supplementary Table 1). Embryos are then rinsed with PTW and refixed with Formalin 10%. Then, embryos are washed with PTW 4 times for 5 minutes. Then, embryos are incubated with *hybridization buffer+* 2h at 65°C. Next, in another recipient, probes are denatured by incubation at 68°C for 5 minutes to denature the probes and then incubation in ice for 2 minutes. Finally, denatured probes are added to embryos and incubated overnight at 65°C.

#### Day 2: *Post hybridization*

Probes are recovered. Embryos are washed 3 times for 30 minutes in *Hybridization Buffer–* at 65ºC. Embryos are washed for 15 minutes in 2X SSC at 65°C. Embryos are washed 2 times for 15 minutes in 0.2X SSC at 65ºC. Next, embryos are washed 3 times for 15 minutes in PTW at RT. Next, embryos are blocked in blocking buffer (PTW+10%FBS) for 1h at RT. Next, anti-digoxin antibody is added at 1:1000 and incubated O/N at 4°C.

#### Day 3: *Washing and Colour development*

Embryos are washed with 2 quick rinses in PTW. Next, embryos are washed 4 times for 15 minutes in PTW. Next, embryos are washed 3 times for 5 minutes with freshly prepared AP buffer. Next, embryos are incubated in 1*µ*L of NBT and 3.5*µ*L per mL of AP buffer. Reaction is allowed to develop for the same amount of time for each group. When colour is fully developed, embryos are washed 3 times for 10 minutes each with PTW. Finally, embryos are transferred to increasing concentrations of Glycerol (30%-50%-70%-100%).

Buffers: Hybridization buffer +: 50% Formamide, 25% SSC 20x pH=7 in H2O miliQ, 0,3% Heparin 50mg/microL; final conc = 150mg/mL, Torula RNA 5mg/mL, 1% of Tween 10%, H2O miliQ up to final volume

Hybridization buffer −: 50% Formamide,10% SSC pH= 7, 1% Tween 10%, H2O miliQ up to final volume.

AP Buffer: 2% NaCl 5M, 10% Tris pH 9,5 1M 5% Mg Cl2 1M, 10% Tween 10%, H2O miliQ up to final volume.

SSC 2X: 10% SSC pH7, 20X 1% Tween 10%, H2O miliQ up to final volume.

### D. Sample mounting, image acquisition and processing

Samples are mounted in homemade chambers with glass slides using 25-50 *µ*L of RapiClear© 1.49 (SunJin Lab). Despite the high transparency of the zebrafish embryo at these stages, the addition of a clearing agent is highly important to ensure sufficient resolution in the vertical planes, which are the ones that limit 3D automated segmentation in confocal reconstructions (18).

Confocal images are captured using a Leica SM800 confocal microscope, with a thickness of 1 *µ*m determined by the pinhole settings. An overlapping of 0.2 *µ*m between confocal sections is established to ensure proper 3D reconstruction. The resolution in the confocal plane is 325×325 *µ*m, while the resolution in the Z direction is determined by the number of confocal sections (ranging from 64 to 120 *µ*m).

Processing and test of computational time was performed in a Personal computer equipped with Windows 11 Pro running on a 13th-generation Intel©Core i7 13/100 2.10 GHz, equipped with 32 GB RAM.

Image processing is conducted using Julia (98) scripts and packages, Python, and FIJI macros (99). Isolation of the neural retina in the image is achieved using a custom macro designed to define the bounding boxes of the region of interest in the XY and XZ planes in To-Pro3 nuclear staining images. The lens is extracted by approximating it to an ellipsoid using the *3D Draw Shape* plugin (100). Further details regarding specific filtering and image processing for each channel can be found in the Supplementary Information.

Automated segmentation was performed using OSCAR (Object Stitching by Clustering of Adjacent Regions) (17), designed based on Machine Learning, dimensionality reduction, nonlinear fitting, and statistical algorithms to bypass potential segmentation errors caused by a poor signal-to-noise ratio (typical of images acquired *in toto* or *in vivo*). We have shown previously that OSCAR shows higher accuracy (based on F1 score, IoU, and Friedman-Rafsky score) than other commonly used tools for 3D segmentation, while requiring less computational resources. OSCAR is freely available in the GitHub repository https://github.com/davidgmiguez/OSCAR.

### E. Automated identification of cell identity based on double staining

Since variability in Atho7 levels is high, even between cells in the same sample, identifying cells as cycling progenitors or terminally differentiated using a single threshold becomes unreliable. Alternatively, we designed a strategy based on levels of Atoh7 and Sox2, which are highly complementary. To do this, we have updated our previously published tool OSCAR (17) to establish cell identity as *P, D*, or *IP* based on Atoh7 and Sox2 levels. A major advantage of this approach is that the thresholds can be defined pragmatically and not based on arbitrary manual settings. The algorithm works as follows:

a. First, the algorithm normalizes the intensity of each confocal plane of each channel, containing information relative to each staining.
b. Next, the total intensity level inside each object in 3D (3D-Projection in the nomenclature used in (17)) is computed for each channel.
c. Depending on the particular characteristics of each staining, a threshold is applied to define cells positive for each staining (for instance, GFP levels in differentiated cells keep accumulating, resulting in much higher levels than recently differentiated cells).
d. The identity of each 3D-Projection is then decided by direct comparison of these two values. In this case, cells with intensity *I*_*Sox*2_ *>* 2*× I*_*GFP*_ are identified as Progenitors, cells with intensity *I*_*GFP*_ *>* 2 *× I*_*Sox*2_ are identified as Differentiated, and cells with intensity 1*/*2 *< I*_*Sox*2_*/I*_*GFP*_ *<* 2 are identified as Intermediate Progenitors (29).

### F. Generation of output figure

Digital reconstructions generated from OSCAR’s output are generated using “3D Viewer” and “3D Draw Shape”, within the 3D ImageJ Suite package in FIJI (100, 101). This way, 3D-Projections are depicted as ellipsoids positioned at their centroid, with orientation and dimensions determined by their x-length, y-length, and z-length. Dimensions of the axes are estimated using the median value of the axes of all the 2D-Projections that are part of the 3D-Projection. The Lookup Table (LUT) for each image is adjusted accordingly, for visualization purposes. In the output images of the zebrafish retina, objects are color-coded based on cell identity.

### G. Branching Process tool

Average cell cycle length and mode of division are measured based on a theoretical framework developed by our lab using a branching process formalism continuous in time (35, 86). In brief, this set of equations correlates the temporal evolution in the change of progenitors and differentiated cells in a population of cells with the changes in the average cell cycle length of the population *T* and the average difference between the rate of symmetric proliferative divisions *pp* and symmetric differentiative divisions *dd*. In this general form, the model aims to distinguish if cells are generated from asymmetric *pd* or symmetric (*pp* or *dd*) divisions (these two scenarios are mathematically equivalent). The mathematical formulation allows us to decouple the two main parameters: average mode *pp* − *dd* and average cell cycle *T* (we have shown previously that these two parameters fully characterize the proliferation and differentiation dynamics of the population (35)). This separation allows us to measure *pp* − *dd* and *T* based only on the quantification of the number of cells, using the two equations reproduced below:

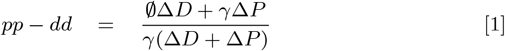

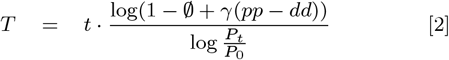

where *pp* − *dd* can be identified with a measure of the differentiation dynamics, and goes from 1 (all divisions being symmetric proliferative) to −1 (all divisions being symmetric differentiative). *T* corresponds to the average cell cycle length of the progenitors. Input values are: *P*_*t*_ and *D*_*t*_ represent the average number of progenitors and differentiated cells at time *t*, while *P*_0_ and *D*_0_ represent the average number of progenitors and differentiated cells at an initial time *t* = 0. Therefore, Δ*P* = *P*_*t*_ − *P*_0_ and Δ*D* = *D*_*t*_ − *D*_0_ represent the change in progenitors and differentiated cells in a window of time Δ*t* = *t* − *t*_0_; *∅* corresponds to the rate of cell death of the progenitors (apoptosis of differentiated cells is not considered, since the geometric series that results for the branching of the Differentiated cells does not converge when apoptosis is considered). Finally, the parameter *γ* defines the growth fraction (rate of progenitors that are actively cycling). *∅* has been estimated using Active Caspase 3 immunostaining for all conditions (Supplementary Figure 2), showing no change with respect to the control. *γ* has been estimated using Pcna immunostaining (Figure 2B). The temporal dependence has been estimated using a spline interpolation with three time points. One of the main advantages of Pcna over BrdU analogs to estimate the growth fraction is that it does not require cumulative curve estimation or previous manipulation of the embryos (it is well known that thymidine analogs affect cell cycle progression (7)). On the other hand, Pcna is down-regulated during M-phase, so it is excluded when estimating the growth fraction, based on pHH3+ staining.

### H. Nonlinear Fitting, Error Propagation and Statistical Analysis

Exponential fitting was performed using the package LsqFit.jl package in Julia, which relies on the Levenberg-Marquardt algorithm as its primary solver for nonlinear least-squares curve fitting.

To facilitate visualization of dynamics and differences between control and experimental conditions, data are accompanied by fitted parametric curves. The best fitting estimation for each condition is determined using Akaike’s Information Criterion (*AIC*). In essence, AIC is used to select the model that provides the best fit to the data while penalizing the inclusion of unnecessary parameters, and is computed as (102):

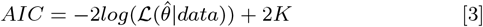

that takes the logarithm of the maximum of the likelihood function of the model, and *K* represents the number of free parameters. Once individual values of *AIC* for each model are computed, the values are rescaled to allow comparison.

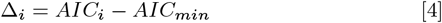

This way, each Δ_*i*_ can be interpreted as the information loss if the model with the minimum *AIC* is not selected. From these values, we can then calculate the likelihood of each model given the data, with the simple transformation *exp*(−Δ_*i*_*/*2) for each of the models. Finally, these likelihood values are normalized and treated as probabilities, as stated in (103):

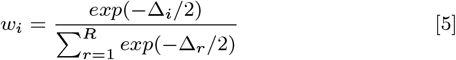

These Akaike weights are considered as the “weights of evidence” in favor of each model as being the best under Kullback-Leibler information theory (a simple metric to measure how far a distribution is from another). Therefore, the model with the highest *w*_*i*_ score can be interpreted to be the most probable model, given the set of data and considered models.

Fitting of experimental conditions is conducted with consideration for mild inhibition levels, resulting in gradual changes in dynamics. Moreover, since inhibitors are introduced at t=20 HPF, values at this time point are equivalent between control and experimental conditions. Thus, the dynamics in experimental conditions are fitted with a single parameter that modulates the control fitting. This approach yields curves for both control and experimental conditions that overlap at 20 HPF and gradually diverge as the drug’s effects manifest over time. These curves serve as input for the branching equations 1-2.

### I. Delta Method for error propagation in nonlinear fitting

The Delta Method (104) is a standard statistical technique used to approximate the variance, and thus the confidence interval, of a nonlinear function of estimated parameters. It is necessary in this context because both our empirical fitting curves and the branching equations are nonlinear functions. The Delta Method works by approximating the variance and standard errors, using the property that the margins of any regression (i.e., values that interpret the effects of each variable) are typically infinitely differentiable functions. This method employs a first-order Taylor series expansion of the inverse link function of the regression to approximate the margin in the vicinity of the data, accomplished through three main steps:

- Calculate the Jacobian matrix of the inverse link function of the fitting.
- Compute the variance-covariance matrix
- pre- and post-multiply the partial derivatives of the inverse link function by the original variance-covariance matrix from the regression.

Here, we use the Delta Method to obtain an estimation of the confidence interval *CI* of the nonlinear fitting of *P* and *D* (i.e., a 95% *CI* marks the range where the true fit lies with 95% confidence interval). These *CI* are illustrated as ribbons in the plots of *P* and *D*. This way, when the ribbons between two regressions do not overlap, we can assume that the two regressions (including the ones with different functional forms) are different with statistical significance.

Values of *P* and *D* selected at random inside these confidence intervals using a Monte-Carlo algorithm are then used as input of the branching equations to obtain the corresponding *CI* in the predictions of *pp* − *dd* and *T* at each time point (35). This prediction is built by recursively adding to the previous value of *P* and *D* cells, a random number drawn from a Gaussian distribution with mean equal to the increment predicted by the fitting, and variance corresponding to the one calculated with the Delta method in *P* and *D*.

This way, the *CI* of the predictions for *pp* − *dd* and *T* (represented as ribbons in the plots) illustrate the uncertainty in the prediction for *pp* − *dd* and *T* (higher dispersion in the experimental data for *P* and *D* results in larger *CI* in the fitting of *P* (*t*) and *D*(*t*), that in turn results in higher uncertainty in the value of *pp* − *dd* and *T* predicted by the branching).

Although the Delta Method is not perfect, it provides several advantages. First, the *CI* is different in different regions of the regression, meaning that two curves can be statistically similar in one region but different with statistical significance in other regions (as in our case, where all curves start at the same point and gradually separate as time progresses).

Finally, the confidence intervals from the Delta Method are propagated using Monte Carlo simulations (105) to assess the uncertainty in the calculation of *pp* − *dd* and *T* (detailed explanation provided in Supplementary Information).

### J. 3D spatial point-pattern clustering analysis

Spatial point-pattern analysis was performed on the centroids provided by the OSCAR software (17), which represent the spatial coordinates of segmented objects within each 3D volume. Object labels were used to distinguish groups for intra- and inter-cluster analyses. The spatial organization was quantified using the three-dimensional form of Besag’s *L* function (106), computed as the variance-stabilized transformation of Ripley’s *K* function (107, 108), and plotted as *L*(*r*) − *r*, (positive values indicate clustering and negative values indicate dispersion relative to complete spatial randomness (CSR)).

Centroid, radius, and rotation matrix were obtained by performing an eigen-decomposition of the covariance matrix of the centroid coordinates, yielding the principal axes and scaling factor required to fully enclose the dataset. For each point, the shortest distance to the ellipsoid boundary was computed analytically in the aligned coordinate system using

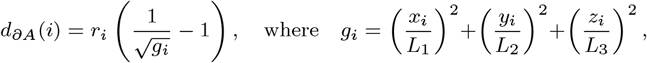

and (*L*_1_, *L*_2_, *L*_3_) are the ellipsoid semi-axes. This distance was used to compute an isotropic edge-correction weight for each reference as point *i* and distance *r*:

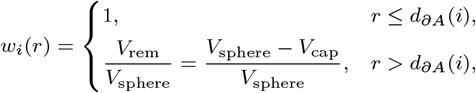

where 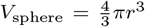 and 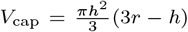 is the volume of the spherical cap^3^truncated by the e^3^llipsoid boundary at height 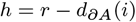.

The unbiased estimator of the three-dimensional *K* function was then defined

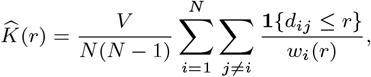

where *V* is the ellipsoid volume and *d*_*ij*_ the Euclidean distance between points *i* and *j*. The transformed *L* function was obtained as

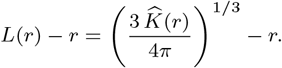

For bivariate analyses, where spatial relationships between two point sets were assessed (e.g., between distinct OSCAR-defined clusters), the estimator was adapted to the cross-type form

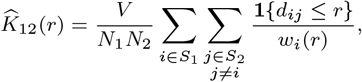

with *S*_1_ and *S*_2_ denoting the reference and neighbor point sets, respectively.

Monte Carlo simulations were used to generate CSR reference envelopes within the same ellipsoidal boundary, employing random uniform point generation in the ellipsoid’s principal-axis coordinate system. A total of 99 simulations were performed, and the 2.5% and 97.5% quantiles of simulated *L*(*r*) − *r* values defined the 95% confidence envelopes. Observed deviations of the empirical *L*(*r*) − *r* curve beyond these envelopes were interpreted as statistically significant clustering or dispersion at the corresponding spatial scales.

### K. Agent-Based Modeling of proliferating and differentiating population

The cell population dynamics were simulated using a discrete-time, age-structured stochastic model implemented in Julia, and designed to allow for multiple independent runs (simulations) to capture the intrinsic variability inherent to the stochastic process. The simulation tracked two distinct populations: progenitor cells (*P*) and differentiated cells (*D*).

To model asynchronous cell cycling, each progenitor cell was assigned an initial age (*A*_*i*_) sampled from a uniform distribution between 0 and *< T >*, where *< T >* corresponds to the mean cell cycle time, defined by the user. To simulate variability in the length of the cell cycle between the progenitors, the value is sampled from a set of randomly distributed discrete values between *< T >* − 2 and *< T >* +2 hours.

At the starting point of our simulations, we set an initial number of *P*_0_ progenitor and differentiated *D*_0_ cells; the simulation progresses in discrete time steps (Δ*t* = 1 hours). Within each step, the following sequence of events occurs:

- **Age Progression:** The age of every existing progenitor cell is incremented by 1 (*A*_*i*_ ← *A*_*i*_ + 1).
- **Division Check:** A progenitor cell is marked for division if its age reaches or exceeds the cell cycle time *A*_*i*_ ≥ *T*.
- **Division Outcome:** Upon division, the mother cell is removed, and the fate of the two resulting daughter cells is determined probabilistically by the parameter *pp* − *dd* ∈ [ 1, − 1]. This parameter governs the balance between self-renewal and terminal differentiation:
- The probability of producing **two new progenitor cells** (self-renewal) is: 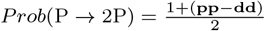
- The probability of producing **two differentiated cells** is: 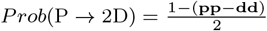
- **Post-Division State:**
- If the division results in two new progenitors, their ages are both reset to **0**, and they are added to the progenitor population.
- If the division results in two differentiated cells, the total count of *D* cells is incremented by two. Differentiated cells are considered non-cycling and accumulate throughout the simulation without further division.

The simulation was run for a total of 28 time steps (to mimic the 28 hours of the experimental data). 10 independent simulations were performed for each condition.

## L. Code Availability

The code generated to run the numerical model that is part of the present manuscript and reproduce the numerical results (109) is available at https://github.com/davidgmiguez/AgentBase.git and https://doi.org/10.5281/zenodo.21709014. OSCAR is freely available in the GitHub repository https://github.com/davidgmiguez/OSCAR and https://doi.org/10.5281/zenodo.21709029.

## M. Data Availability

Data used for the plots and the analysis is available in https://doi.org/10.6084/m9.figshare.33126719. Data format is available as summary files in txt format, organized as columns.

## O. Author contribution

D. P-D, M. L-T, and D. G. M. designed the experiments, D. P-D, M. L-T, D. M-D. G. N-M and D. G. M. performed the experiments and analyzed the data, D. G. M. designed the computational model, performed the simulations, analyzed the results, provided funding and wrote the manuscript.

## ACKNOWLEDGMENTS

We acknowledge all CBMSO (Centre for Molecular Biology Severo Ochoa, CSIC–UAM) facilities. We thank Pablo Martinez-Martinez (Universidad Autonoma de Madrid) for help developing the Delta Method for error propagation. MLT, DPD, and DGM designed research. MLT and DPD performed research. MLT, DPD, GNM, DMD and DGM analyzed results and DGM wrote the manuscript. Authors declare no competing interests. DPD (Author One) contributed equally to this work with MLT (Author Two).

