## Supplementary material for "Exponential growth and neuronal yield in the vertebrate developing retina are mediated by Hedgehog signaling": Suplementary data

July 31, 2026

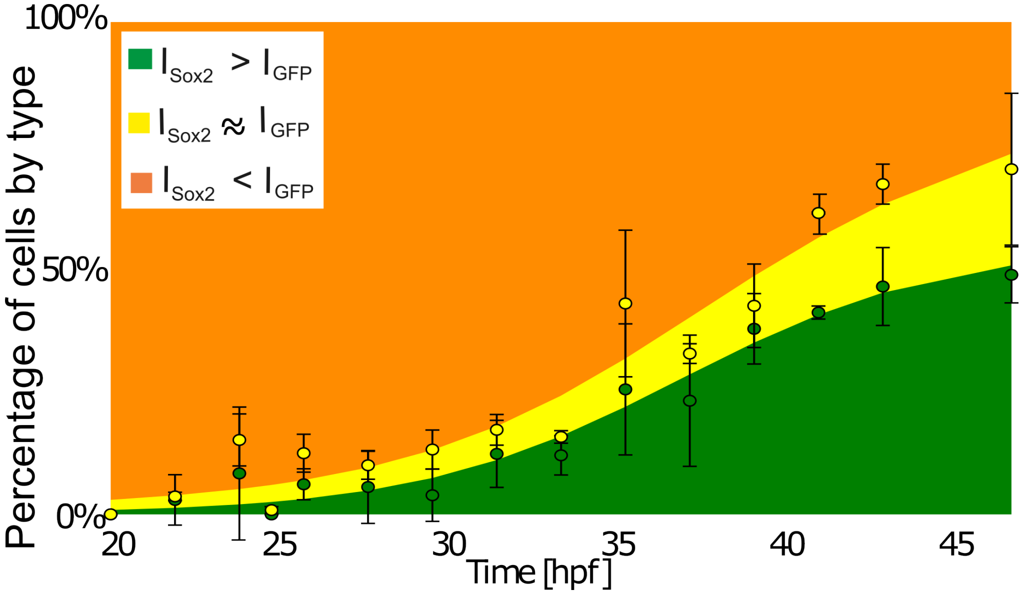


Figure 1: *

**Supplementary Figure 1: Distribution of the three cell types at different HPF**. $P$ depicted as red, $D$ as green, and $IP$ as yellow. The percentage of $D$ and $IP$ increases as development progresses.

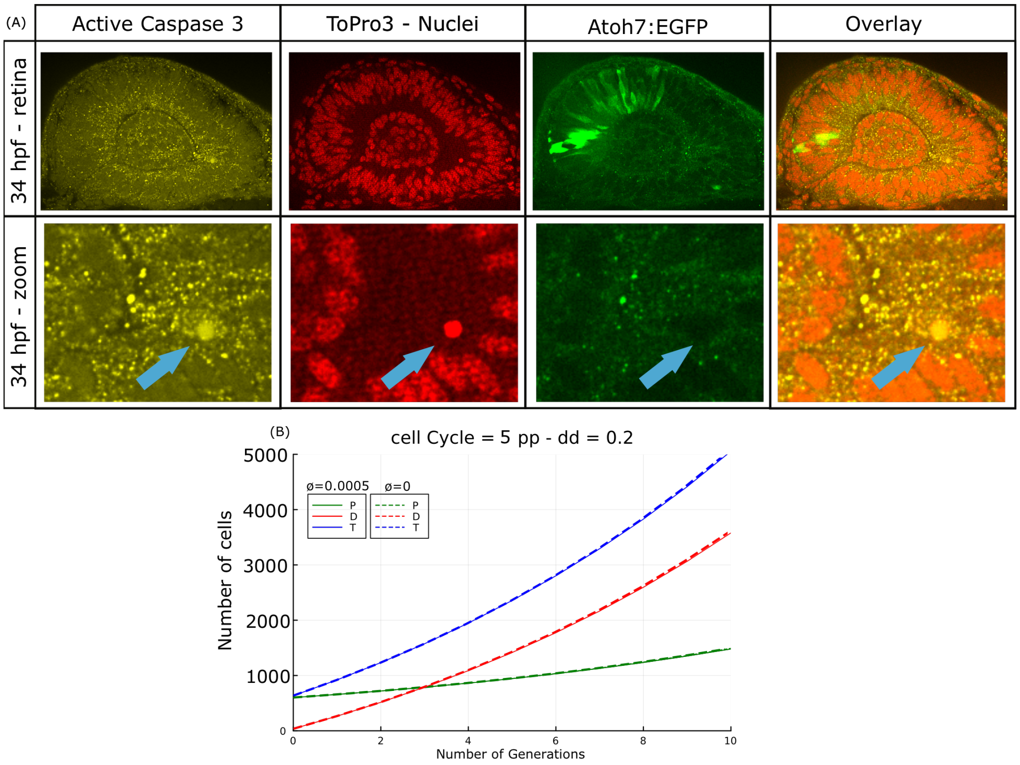


Figure 2: *

**Supplementary Figure 2: Positive control for immunostaining against Active Caspase 3.** (A) Confocal section of a zebrafish retina at 34 HPF imaged *in toto* as positive control showing cells positive for immunostaining against Active Caspase 3. (B) Comparison of numerical simulations of the agent-base model for proliferating and differentiating populations of cells, with (solid lines) and without (dashed lines) apoptosis.

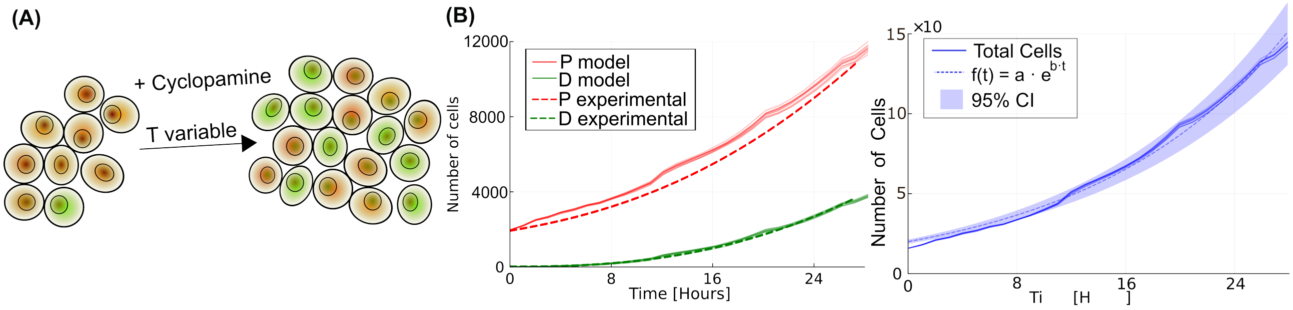


Figure 3: *

**Supplementary Figure 3: Plot of the model prediction for conditions of Cyclopamine-treated embryos.** (A) Scheme of the growth of the population of progenitors and differentiated cells. (B) The numerical simulations (solid lines) using as input the values of *pp-dd* and $T$ predicted by the branching equations recover the experimental data (dashed lines). There is a higher discrepancy in Cyclopamine conditions compared to the control (Figure 5A of the main text), probably due to a change in the growth fraction or the apoptosis rate in conditions of cyclopamine. The systems still grow in terms of total number of cells (blue lines, right panel) following a clear exponential profile. Ribbons represent the 95% confidence interval of the fitting.

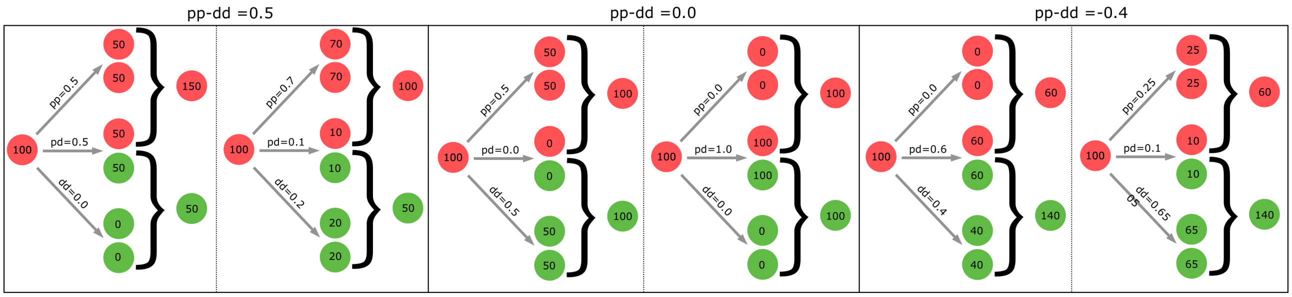


Figure 4: *

**Supplementary Figure 4: Number of Progenitors (red) and Differentiated cells (green) in the context of the probabilities for the three different modes of division (pp, pd, and dd).** Three conditions are illustrated (increase, homeostasis, and decrease in progenitors). For each condition, two scenarios are illustrated with different probabilities for each mode of division, but maintaining the same value of $pp - dd$ (remember that in the branching framework, the rates are probabilities, so pp+pd+dd=1). We can see that after one iteration (discrete time), the final number of P and D is defined by the difference between the rates of symmetric proliferative $pp$ and symmetric differentiative $dd$, so the framework cannot be used to measure the individual rates.

Figure 5: *

**Supplementary Movie 1:** 3D view of the input and output images of the zebrafish retina corresponding to Figure 3B.

Figure 6: *

**Supplementary Movie 2:** Confocal planes of the input and output images of the zebrafish retina corresponding to Figure 3B.

| Stage (HPF) | Incubation time(min) |
|---|---|
| 20 | 10 |
| 22 | 12 |
| 24 | 14 |
| 26 | 16 |
| 28 | 18 |
| 30 | 20 |
| 32 | 23 |
| 34 | 25 |
| 36 | 28 |
| 38 | 31 |
| 40 | 33 |
| 42 | 36 |
| 44 | 39 |
| 46 | 42 |
| 48 | 45 |

Table 1: *

Supplementary Table 1: Proteinase K exposure times for different stages.

| Figure | Condition | $pp - dd(t)$ | $T(t)$ [hours] | $\gamma(t)$ |
|---|---|---|---|---|
| 1B | High Proliferation | Constant: 0.95 | Constant: 6 | 1.0 |
| 1C | Homeostasis | Constant: 0.0 | Constant: 6 | 1.0 |
| 1D | Gradual Shift | Linear (1.0 to -0.5) | Constant: 6 | 1.0 |
| 1E | Sharp Switch | 1.0 for $t < 14$h, -0.5 for $t \geq 14$h | Constant: 6 | 1.0 |
| 5A | Variable T | From 4D (blue line) | From 4D (yellow line) | spline |
| 5B | Constant T | From 4D (blue line) | Constant: 8 | spline |
| SuppFig 3 | +Cyclo | From 6F (solid blue) | From 6G (solid yellow) | spline |

Table 2: *
Supplementary Table 2: Parameter values of the model used for the figures.

Table 3: Fitting results for numerical simulations and experimental data.

| Data input for fitting | Condition | A | | B | |
|---|---|---|---|---|---|
| | | mean | 95% CI | mean | 95% CI |
| Numerical simulations | (Figure 1b) pp-dd=0.95 | 2191 | 68 | 0,068 | 0,001 |
| | (Figure 1c) pp-dd=0.05 | 2617 | 250 | 0,042 | 0,005 |
| | (Figure 1d) pp-dd variable (smooth) | 2419 | 245 | 0,061 | 0,004 |
| | (Figure 1e) pp-dd variable (sharp) | 2447 | 312 | 0,060 | 0,005 |
| Experimental data | (Figure 5a) Fixed T | 2343 | 202 | 0,069 | 0,004 |
| | (Figure 5b) Variable T | 2006 | 141 | 0,077 | 0,003 |

Table 4: *
Supplementary Table 3: Parameters for exponential fitting $N = a \cdot e^{b \cdot t}$ for the number of total cells in numerical simulations and experimental data.
